# Understanding neural circuit principles for representation learning through joint-embedding predictive architectures

**DOI:** 10.1101/2025.11.25.690220

**Authors:** Ashena Gorgan Mohammadi, Manu Srinath Halvagal, Friedemann Zenke

## Abstract

Whether tracking prey or evading a predator, an animal’s survival depends on both identifying objects in the environment and anticipating how they will move. To guide behavior, the brain must therefore extract information about object identities and their dynamics from entangled sensory inputs. How it accomplishes this feat remains an open question. Classical predictive coding theories propose that this ability arises by comparing predicted sensory signals with actual inputs and reducing the associated prediction errors. While such models capture important aspects of cortical computation, they typically focus on faithfully predicting sensory input and do not explicitly address how abstract, untangled representations of objects and their dynamics emerge solely through experience. Here, we develop a theory of representation learning in neural circuits that shifts the focus from prediction in the input space to prediction in representation space, without relying on external supervision or labeled data. Specifically, we introduce recurrent predictive learning (RPL), a recurrent joint-embedding predictive architecture inspired by self-supervised learning, that learns abstract representations of object identity and their dynamics and predicts future object motion from continuous sensory streams. While local learning rules alone can learn invariant object identity, we find that capturing object dynamics requires a circuit that explicitly generates and minimizes internal prediction-error signals. Crucially, the model learns sequence representations that resemble successor-like representations observed in the human primary visual cortex. The model also develops abstract sequence representations comparable to those reported in the macaque prefrontal cortex. Finally, we demonstrate a hierarchical version of RPL and discuss possible mappings onto cortical microcircuits. Our work establishes a circuit-centric theory framework that provides new perspectives on how the brain may acquire an internal model of the world through experience.

## Introduction

It takes us a split second to recognize an oncoming car at an intersection, to anticipate the trajectory of a ball thrown to us, or to realize that we are being chased by a dog in the park. Importantly, we have no difficulty assessing these situations and responding to them, despite never having seen this particular car, ball, or dog. To accomplish this feat, our nervous system faces a twofold challenge. On the one hand, it must detect abstract patterns from sensory input and form invariant object representations, allowing us to recognize, for instance, a dog. This task is hindered by the fact that distinct sensory stimuli are often entangled, requiring complex hierarchical networks to untangle them (***Figure 1***). On the other hand, our brain must also infer equivariant features of the dog’s dynamics, such as its position, velocity, and angle of attack. After all, our actions depend on whether it is moving toward or away from us. This can be achieved by learning structured representations of the dynamic latent variables, such that temporal trajectories in the latent space are internally represented by corresponding low-dimensional trajectories. While a plethora of experiments have shown that sensory cortices encode this information (***Logothetis and Sheinberg, 1996***; ***Hubel and Wiesel, 1962***; ***Duhamel et al., 1992***; ***Whitney et al., 2003***; ***Johnson et al., 2023***; ***Maunsell and Van Essen, 1983***; ***Hennestad et al., 2021***; ***Priebe et al., 2006***; ***Roitman et al., 2005***), the mechanisms underlying the plasticity that forms these representations remain elusive. Previous modeling work indicated that minimal extensions to local Hebbian learning rules can learn invariant object representations in deep neural networks (***Illing et al., 2021***; ***Halvagal and Zenke, 2023***). However, it remains unclear whether similar mechanisms also yield equivariant, or structured, representations of the dynamic latent variables. Yet, both invariant *and* equivariant representations are necessary to respond appropriately in novel situations.

**Figure 1.**
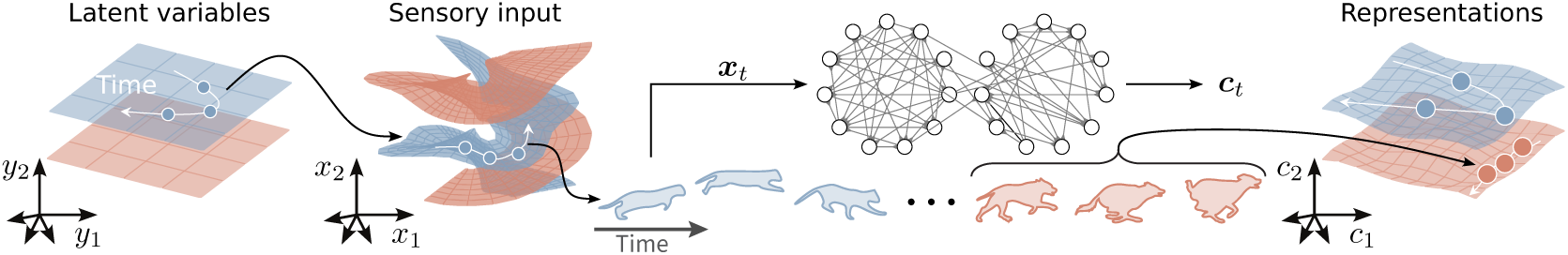
Extracting latent variables from sensory stimuli. Animals and humans must infer latent variables ***y*** such as object identity, orientation, and velocity from ambiguous and noisy sensory stimuli. This task is computationally demanding because sensory stimuli ***x*** are typically entangled (left). Untangling these input manifolds to approximate the underlying latent variables ***c****_t_* (right) requires multi-layer networks (middle). How our brains achieve this untangling remains a central question in neuroscience.

Another influential idea of how representation learning works in the cortex is predictive processing (***Keller and Mrsic-Flogel, 2018***), typified by classical predictive coding models (***Rao and Ballard, 1999***). These propose that the brain acquires an internal model of the world that enables it to anticipate incoming sensory input. In this framework, cortical networks generate predictions of expected sensory stimuli and compare them to the actual input, yielding prediction errors. Minimizing these errors supports perceptual inference of latent variables, representation learning, and the continual refinement of the internal model (***Rao and Ballard, 1999***; ***Singer et al., 2018***; ***Lotter et al., 2020***; ***Hertäg and Sprekeler, 2020***; ***Recanatesi et al., 2021***; ***Mikulasch et al., 2023***; ***Levenstein et al., 2024***; ***Nejad et al., 2025***; ***Rao, 2024***; ***Taylor et al., 2024***).

While a compelling idea, classical predictive coding models have one major drawback: they are generative models that construct predictions in the input space. Consequently, such models require a decoder network that predicts, for example, the pixel-level image of an incoming dog, which is then compared to the actual sensory experience, rather than comparing internal representations directly. This subtle distinction has profound consequences for the neural circuit architecture required, specifically where in the circuit prediction errors are computed, how they interact with representation neurons, and how they guide learning.

A growing body of work has shown that classical predictive coding models trained as fully generative systems often struggle to yield representations that support abstraction, invariance, or object-level discrimination. In particular, models that minimize local prediction errors greedily at each hierarchical stage tend to learn representations that remain closely tied to input statistics, limiting their usefulness for downstream classification or invariant recognition (***Tscshantz et al., 2023***; ***Oliviers et al., 2025***). This limitation is also evident in large-scale models such as PredNet (***Lotter et al., 2020***). While PredNet achieves its best performance when prediction errors are optimized only at the input (pixel) level through end-to-end training, thereby allowing deeper representations to be shaped indirectly, its representational and predictive performance degrades when prediction errors are minimized across all hierarchical levels. Together, these findings suggest a tension between generative, reconstruction-driven objectives and the learning of abstract representations, motivating predictive frameworks that operate directly in the representation space rather than the sensory domain.

Finally, classical predictive coding posits that predicted sensory information is represented with reduced neural activity in early sensory cortices due to suppressive top-down feedback. However, a growing body of neurophysiological evidence indicates that top-down input from higher cortical areas can enhance representations of behaviorally relevant or attended stimuli (***Westerberg and Roelfsema, 2025***). Consequently, whether and how we can reconcile predictive processing with cortical anatomy, abstract representation learning, and empirical observations of inter-areal interactions remains a central outstanding question in circuit neuroscience (***Keller and Mrsic-Flogel, 2018***; ***Solomon et al., 2021***; ***Gillon et al., 2024***; ***Richter et al., 2024***; ***Furutachi et al., 2024***; ***Westerberg and Roelfsema, 2025***; ***Vasilevskaya and Keller, 2026***).

A natural alternative is to abandon explicit generative modeling and instead compute prediction errors entirely within internal representation space. This strategy has proven effective in recent self-supervised learning frameworks (***Balestriero et al., 2023***). By eliminating the need for a decoder that reconstructs sensory input, such an approach would avoid the difficulties associated with generative prediction in the input space. However, representation space prediction introduces a new challenge: *representational collapse*, wherein networks converge to trivial solutions, e.g., constant representations, that are perfectly predictable but devoid of useful information. Preventing collapse requires additional constraints, such as variance regularization techniques (***Bardes et al., 2022***; ***Halvagal and Zenke, 2023***; ***Niu et al., 2024***), negative samples (***Oord et al., 2018***; ***Illing et al., 2021***), or a joint-embedding predictive architecture (JEPA) (***LeCun, 2022***).

JEPAs avoid collapse by computing prediction errors between internal representations using a dedicated predictor network (***Grill et al., 2020***; ***Chen and He, 2021***; ***Assran et al., 2023***; ***Ghaemi et al., 2024***; ***Bardes et al., 2024***; ***Assran et al., 2025***; ***Balestriero and LeCun, 2025***). Beyond biological plausibility, this strategy offers another distinct advantage: by dispensing with reconstruction objectives, the network can devote its coding capacity to abstract, behaviorally relevant variables such as object identity and future dynamics, e.g, “the dog behind me is about to pounce,” rather than low-level sensory details that may be incidental or task-irrelevant, such as the particular sheen of the dog’s fur around the ears (***Balestriero and LeCun, 2024***; ***Littwin et al., 2024***).

Despite these advantages, most JEPAs developed in machine learning are poor models of cortical computation. They rely on architectural elements with limited biological grounding, such as transformers, and training strategies, such as token masking, while focusing mainly on static images, hand-crafted transformations, and invariant representations geared towards object classification. As a result, it remains unclear whether a JEPA broadly consistent with cortical circuit structure and function exists. A minimal candidate must instead rely on recurrent neural dynamics, learn from streaming sensory input without masking, support both invariant and equivariant representations, and reproduce key neurophysiological observations.

Here, we introduce such a model. Our goal is to develop a circuit-level theory of representation learning that explains how abstract representations could emerge in cortical networks and to identify qualitative correspondences and testable predictions, rather than to quantitatively fit neural data. To that end, we propose recurrent predictive learning (RPL). This minimal circuit model learns linearly decodable object representations, including their associated motion variables, from videos of moving objects without requiring any labels. As we will see, it yields abstract representations spanning multiple timescales, consistent with representations in the macaque prefrontal cortex (PFC) (***Bellet et al., 2024***). In addition to representation learning, RPL learns a dynamic world model, which represents future stimuli, similar to human V1 (***Ekman et al., 2023***). To accomplish this, RPL requires distinct feedforward and recurrent components, each playing a dedicated functional role. In a hierarchical setting, prediction errors are computed in local prediction circuits; this organization allows for an interpretation in terms of necessary neuronal circuit elements. In summary, we introduce a circuit-centric network model and learning strategy that extends the notion of predictive processing to internal representation prediction, providing a principled set of components for learning structured representations in the brain.

## Results

To study the putative mechanisms by which sensory networks extract latent variables, such as invariant object identity, and equivariant latent variables, such as position, velocity, and orientation, we simulated deep neural network models with a specific structure. Broadly speaking, these networks consist of three functionally distinct components (***Figure 2a***). First, a feedforward “encoder” network nonlinearly encodes sensory stimuli into the intermediate embedding ***z*** with minimal to no history dependence. Second, a recurrent neural network, the “integrator,” successively integrates these embeddings to form the internal representations ***c***. Third, the predictor network *f* takes the representation ***c****_t_* at time *t* to predict the embedding ***z****_t_*_+Δ_ an instant later. Finally, the resulting prediction error

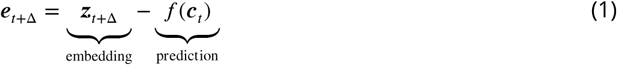

**Figure 2.**
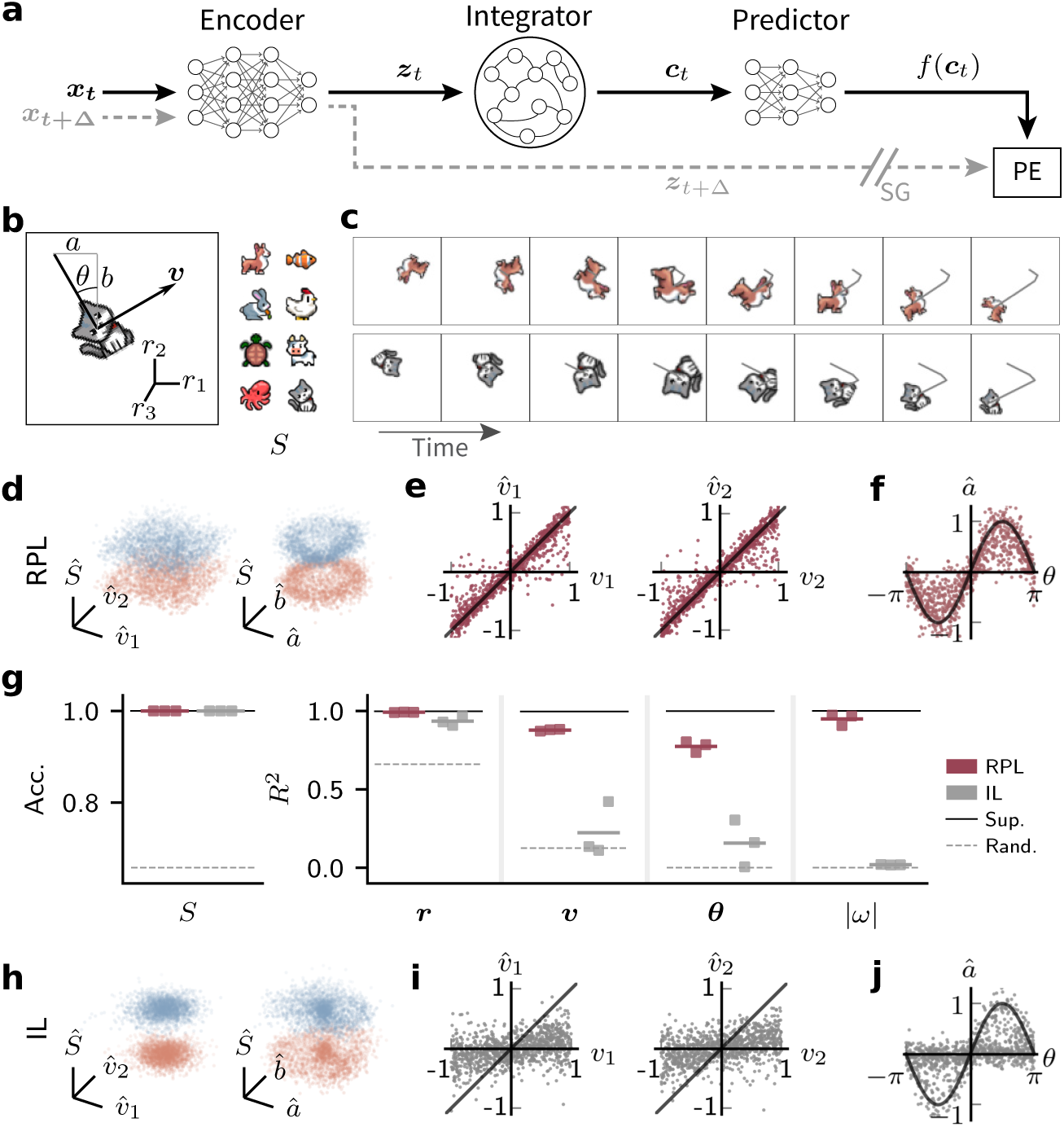
Learning latent variables from motion stimuli without reconstruction and labels. **(a)** Schematic of the network model. RPL consists of different network components. First, the “encoder” encodes sensory stimuli ***x*** into intermediate embeddings ***z***. Subsequently, the “integrator” synthesizes these embeddings into the internal representation ***c***. Finally, the “predictor” network predicts future embeddings ***z****_t_*_+Δ_ from ***c****_t_* yielding the associated prediction error (PE). The learning objective is to minimize this PE. (b) Synthetic data generation paradigm with known latent variables. We generate motion trajectories by moving a 2D animal image, chosen randomly from one of eight categories (right), within a square arena (left). Each frame is characterized by the object position *r*_1∕2_, its scale to mimic position in the *r*_3_ direction, and the orientation angle *θ*. For each trajectory we randomly select a constant velocity ***v*** and angular velocity 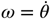. To obtain pixel images that serve as network inputs, we further place these motion trajectories on a gray background and add Gaussian pixel noise, and elastic collisions at the borders (Methods). (c) Two example trajectories shown frame-by-frame. (d) Scatter plots of linearly decoded latent variables from learned internal representations ***c*** for a representative sample of cat (blue) and dog (red) input images. Velocity (*v*_1_, *v*_2_) and object identity *S* (left). Orientation (*a*, *b*) and *S* (right). (e) Decoded velocity components 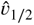 as a function of ground truth on held-out data. (f) Same as (e) but for â as function of *θ*. The model (RPL) has formed untangled and structured representations of the latent variables. (g) Linear readout accuracy of animal categories *S* (left) and coefficient of determination (*R*^2^) (right) for position ***r***, velocity, orientation ***θ*** = (*a*, *b*), and angular speed |*ω*| for linear decoders trained on the internal representations ***c*** for *n* = 3 independent networks, trained with the RPL and invariance learning (IL) objectives. Decoding performance for *n* = 1 supervised end-to-end training (Sup.) and a randomly initialized network (Rand.) are given for reference. RPL decoding accuracy for object identity, and *R*^2^ values are close to the performance of networks trained end-to-end with supervised learning, whereas IL fails to accurately represent the velocity, orientation, and angular speed.(h,i,j) Same as d-f but for IL. ***Figure 2—figure supplement 1***. Sample video frames input to the models. ***Figure 2—figure supplement 2***. Full comparison of representations learned by different objectives. ***Figure 2—figure supplement 3***. 3D projections of learned representations for all trained models. ***Figure 2—figure supplement 4***. Linear projections of learned representations of motion factors. ***Figure 2—figure supplement 5***. Orthogonality of learned representations of motion factors. ***Figure 2—figure supplement 6***. Effect of pixel noise and frame dropout on the learned representations.

serves as an instructive learning signal to optimize the model parameters (Methods). Importantly, this unsupervised learning signal is fully specified in the internal neuronal representation space, localized in time, and can be computed without an explicit prediction or reconstruction of ***x*** in the input space. We collectively refer to network models trained by minimizing the above prediction error as RPL in the following.

To evaluate RPL’s effectiveness in forming untangled representations of both invariant and dynamic variables from object motion sequences, we constructed a synthetic dataset of moving animal images (***Figure 2b***; Methods) with *known* latent variables. To that end, we generated short sequences of one of eight cartoon animal images moving smoothly with a randomly chosen linear and angular velocity within a square arena with elastic collisions at the boundaries (***Figure 2c***). To obtain individual video frames, we sampled the trajectory at discrete intervals, placed each frame in front of a gray background, and added Gaussian pixel noise (***Figure 2—figure Supplement 1***; Methods). Each video frame was characterized by the animal’s identity *S*, its position ***r***, and velocity ***v***, with *r*_1∕2_ defining the planar position and *r*_3_ defining the scale, orientation *θ*, and the corresponding angular velocity *ω*. Finally, to ensure that these latent variables were indeed entangled in stimulus space, we measured how well a linear decoder could predict them from pixel values alone. To that end, we quantified linear classification accuracy and decoding quality (*R*^2^) of dynamic variables. Most variables could not reliably be decoded from the raw input image sequences (***Figure 2—figure Supplement 2***; Supplementary Table S1).

To test whether RPL recovers the underlying invariant and dynamic latent variables, we trained our model on approximately 5.7 h of synthetic videos (Methods). We then visualized the learned representations by plotting two example classes (Cat and Dog) onto their best linear projections of the latent linear velocities *v*_1∕2_. The resulting point clouds faithfully recovered the underlying uniform velocity distribution of the data (***Figure 2d,e***; ***Figure 2—figure Supplement 3a,b***). Similarly, when regressing on *a* = sin *θ* and *b* = cos *θ*, the resulting linear projections revealed an explicit orientation encoding in the internal representations (***Figure 2d,f***). Finally, we quantified classification accuracy and decoding quality of position, velocity, orientation ***θ*** = (*a*, *b*), and angular speed |*ω*|, and saw high decoding performance for all latent variables (***Figure 2g***). The decoding performance of RPL was substantially better than a random untrained baseline model and close to a reference model trained using supervised learning. For comparison to contrastive self-supervised learning (SSL), a powerful machine learning approach that typically violates temporal locality, we also trained and evaluated the same network model using the contrastive predictive coding objective (***Oord et al., 2018***). RPL training yielded better linear decodability of equivariant features (Supplementary Table S1; Methods). Thus, RPL learns untangled representations of object identity and motion variables that can be read out linearly, a computation that downstream neurons can easily implement.

We were curious whether this dual untangling of object identity and motion variables depends on the particular type of JEPA we used, in which the recurrent pathway predicts the feedforward signal. To address this question, we kept the network architecture unchanged and used invariance learning (IL) with a symmetric learning objective similar to previous work (***Bardes et al., 2022***; ***Halvagal and Zenke, 2023***) in which the predictor network predicts its own output instead of the encoder output (Methods). Concretely, this IL objective minimizes the prediction error ***e****_t_*_+Δ_^IL^ = *f*(***c****_t_*_+Δ_) − *f*(***c****_t_*). Networks trained to minimize such IL prediction error formed untangled representations of object identity and position. Still, their representations did not allow reliable decoding of other latent variables such as velocity, orientation, and angular speed (***Figure 2g–j***; ***Figure 2—figure Supplement 3c,d***, ***Figure 2—figure Supplement 4***, ***Figure 2—figure Supplement 5***; Supplementary Table S1). Similarly, optimizing the network using an IL objective at the level of the embedding ***z*** or the representation ***c*** (Eqs. (***4***), (***5***)), resulted in worse decoding performance than the RPL objective (***Figure 2—figure Supplement 2***). Thus RPL’s asymmetric recurrent prediction circuit is essential to learn representations of static and dynamic latent variables and specifically for representing direction of motion or sequence order.

We next wondered whether an alternative asymmetric JEPA implementation, in which the prediction target is the integrator output rather than the encoder output, could also learn to extract both invariant and dynamic latent variables. If so, this would indicate that the asymmetric prediction character of a JEPA is key to learn both invariant and equivariant representations, while being insensitive to where the prediction errors are computed in the circuit, i.e., before or after the integrator. To investigate this question, we defined context-predictive learning (CL) as a third learning paradigm which minimizes prediction errors resulting from predicting the future representations at the level of the integrator ***c****_t_* by minimizing prediction errors ***e****_t_*_+Δ_^CL^ = ***c****_t_*_+Δ_ −*f*(***c****_t_*). We then evaluated the representations using linear probing as before. Again, decoding accuracy from the resulting network representations was worse than RPL, but better than the IL objectives (***Figure 2—figure Supplement 2***). Hence, RPL’s learning objective of predicting embeddings from a context-aware internal representation is crucial for learning untangled representations of invariant and dynamic latent variables.

We speculated that CL’s performance gap may be due to the integrator being able to internally “hallucinate” predictable signals that minimize the prediction error, even though they are unrelated to the input. We reasoned that this issue should become more pronounced when extracting latent variables from noisy input data, thereby requiring the network to rely more heavily on the integrator to infer them by integrating over time. To test this idea, we generated synthetic videos as before while systematically increasing pixel noise and frame dropout, i.e., introducing different numbers of consecutively blank frames at random locations in the video (***Figure 2—figure Supplement 6a***). Consistent with the above hypothesis, we found that the performance gap between RPL and CL widened with increasing noise levels and number of dropped frames (***Figure 2—figure Supplement 6b,c***), while RPL’s overall performance remained relatively stable. Moreover, CL’s performance variability between independently trained models was higher, suggesting that the CL paradigm is less robust. Thus, RPL’s particular asymmetric architecture is essential to robustly untangle dynamic and invariant latent variables. Consequently, we will now focus on comparing RPL to IL for the remainder of the article.

### RPL circuits learn an internal world model

In addition to representing objects and their motion trajectories, animals need to anticipate future states of the world based on past observations. We wondered whether RPL implicitly learns a world model capable of such mental simulation. We reasoned that if the recurrent circuit indeed learns a world model, it should simulate motion trajectories using its internal representations in an autoregressive way. To test this idea, we introduced a dynamic switch in the network setup that routes the predicted embeddings *z̃* back as input to the integrator, replacing the feedforward input from the environment (***Figure 3a***; Methods). Next, we primed a network in feedforward mode with the first eight frames of an animal motion stimulus to allow the model to infer the underlying latent variables. We then switched to feedback mode for 16 time steps. Throughout, we decoded all latent variables from the internal representations as before. We found that the networks in feedback mode simulated motion trajectories that closely resembled the position, scale, and object identity of the actual trajectory (***Figure 3b***; ***Figure 3—figure Supplement 1***). Similarly, the velocity and orientation estimates resembled the ground-truth dynamics (***Figure 3c***). Finally, to check how errors evolved over time, we calculated the average mean squared error over many simulated trajectories for different dynamic variables. This analysis showed that errors dropped rapidly during the inference phase, as expected, and slowly increased during the simulation phase (***Figure 3d***). Together, these findings show that the learned internal representations can reliably simulate future object motion and that RPL learns a world model.

**Figure 3.**
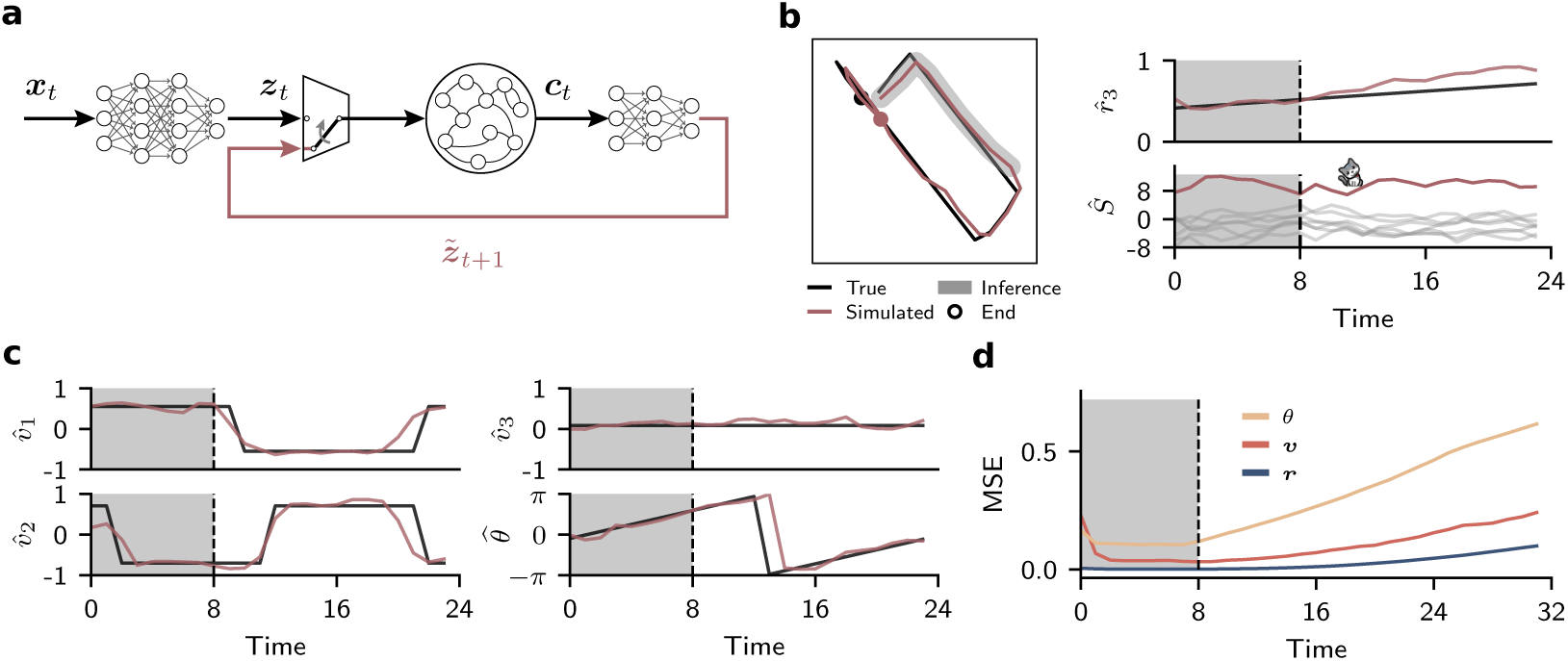
RPL circuits learn an internal world model. **(a)** Setup for simulating trajectories using internal representations. We added a dynamic switch allowing to replace feedforward input to the integrator with internal predictions (cf. ***Figure 1***). To simulate future trajectories, we provide the network with external-world sensory input during an initial inference period. Afterwards, we replaced feedforward input to the integrator with predicted embeddings in an autoregressive manner. **(b)** Example of a decoded trajectory (red) compared to the ground truth (black) in the *r*_1_-*r*_2_ plane (left) and *r*_3_ as a function of time (top right; cf. ***Figure 2a***) as well as the object decoder logits (bottom right). In this example we used an initial inference period of eight frames of a cat trajectory (shaded) followed by 16 steps of internal simulation. **(c)** Decoded velocity and orientation during the inference period (shaded) and simulation. The decoded values closely follow the ground truth. **(d)** Mean-squared error of the decoded quantities compared to ground-truth as a function of time. The error increases monotonically over simulation time. ***Figure 3—figure supplement 1***. More samples of simulated motion trajectories.

### RPL learns variables underlying 3D object motion and lighting conditions

In the moving animal task studied above, the latent-variable dynamics were deterministic and predominantly linear (except at the collision points with the edges), and the frames had a fixed background. Furthermore, each animal category was associated with a single two-dimensional sprite image, reducing the need for representation learning. We wondered to what extent the above findings generalize to more realistic non-deterministic data domains with different objects organized in categories, a larger number of degrees of freedom in object motion, as well as other factors such as changing lighting conditions. Existing benchmark datasets of sufficient complexity for probing equivariance consist of isolated image pairs with a known latent transformation (***Garrido et al., 2023***), which precludes learning from continuous temporal progression of a *sequence* of images that gradually result in a given transformation.

To address this gap, we developed the Temporal Latent Traversal Twelve (TLT12) dataset, building on work by ***Garrido et al. (2023)***. TLT12 consists of 120,000 image sequences with 32 frames each. Each sequence corresponds to a smooth latent traversal of one of 120 objects from 12 categories, including cars, mugs, earphones, and watercrafts, among others (***Figure 4a***; Methods). In contrast to the animals dataset, TLT12 has more factors, greater object diversity, is rendered using true 3D objects under dynamically changing lighting conditions, and the camera position is set at an angle so that the image plane does not trivially align with the coordinate axes (***Figure 4b***). Importantly, the rate of change of the individual factors is not static but changes stochastically following an Ornstein-Uhlenbeck process (Methods). To encourage realistic view angles in the dataset, objects can do full 2*π* rotations about the vertical *r*_3_ axis, whereas rotations about the planar *r*_1∕2_ axes are constrained to lie within a 45% cone. To facilitate future comparison, the dataset comes prepackaged with an 80:10:10 training, validation, and test split.

**Figure 4.**
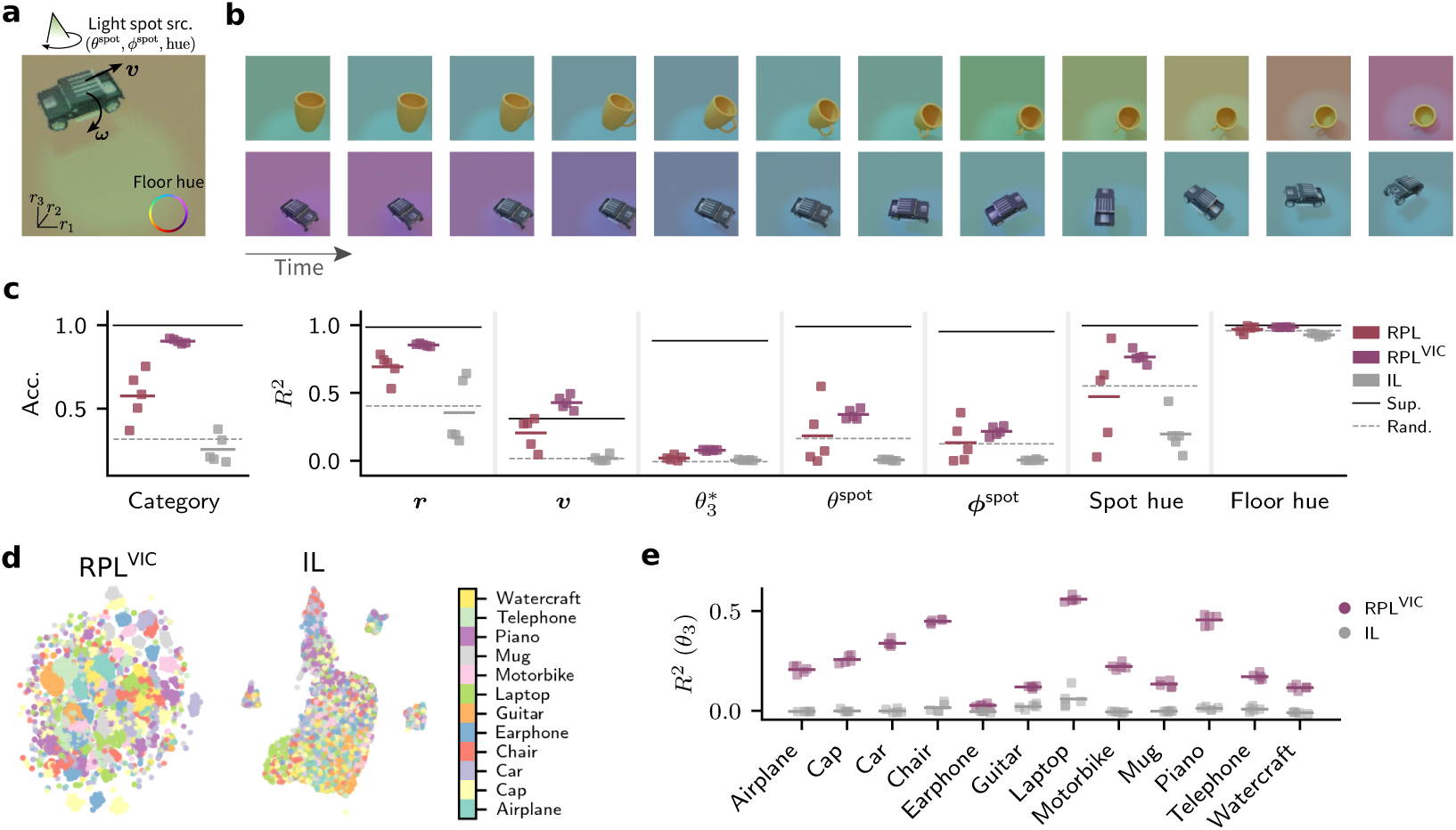
RPL learns variables underlying 3D object motion and lighting conditions. **(a)** Schematic of the latent variables in the TLT12 dataset. The dataset comprises object motion trajectories with changing lighting conditions due to a moving spot and changing floor hue. **(b)** Two example trajectories shown frame-by-frame. Each frame is characterized by the object’s category, position (***r***), velocity (***v***), orientation (***θ***), and angular velocity (***ω***) in 3D space, as well as the spot position characterized by elevation (*θ*^spot^), azimuth (***ϕ***^spot^), and hue plus the floor hue. Crucially, the rate of change of the factors is not constant, but changes itself following an Ornstein-Uhlenbeck process. **(c)** Linear readout accuracy for various latent variables for *n* = 5 independent networks for RPL (red), RPL with VIC regularization (RPL^VIC^) (plum) and IL (gray). Average performance of end-to-end supervised training (black line) as well as untrained networks (dashed line) are given for reference (*n* = 5). RPL consistently outperforms IL. The additional inductive bias in RPL^VIC^ results in more reliable representations of the latent variables. **(d)** Typical examples of two-dimensional UMAP projections of the RPL^VIC^ learned representations (left) and IL (right) colored by object category. RPL^VIC^ representations show more evident categorical organization than IL. **(e)** Linear probing performance (*R*^2^) of per-category linear decoders trained independently. RPL^VIC^ orientation decoding is lower for objects with a bilateral symmetry about the z-direction, e.g., headphones, whereas IL represents orientation poorly for across categories. ***Figure 4—figure supplement 1***. Extended linear decoding performance. ***Figure 4—figure supplement 2***. Linear projections of ground truth and learned representations of translation factors. ***Figure 4—figure supplement 3***. Linear projections of ground truth and learned representations of light spot and floor factors. ***Figure 4—figure supplement 4***. More sample video frames from TLT12. ***Figure 4—figure supplement 5***. Orientation coding in representations learned by RPL and IL models. ***Figure 4—figure supplement 6***. Linear decoding performance on representations of out-of-distribution test samples.

We adapted our network architecture to the larger TLT12 dataset and trained the corresponding networks using RPL as before. Again we used linear probing on the integrator output **c** to assess representational quality. RPL-trained models allowed to linearly decode invariant and dynamic variables (***Figure 4c***; Supplementary Table S2). As expected, IL models performed comparably to random networks or even worse. Notably, IL representations showed a strong bias to represent floor hue, the most salient feature in the dataset, and failed to represent object categories.

Moreover, we noticed that less salient features such as class identity and the light spot position were less consistently encoded across RPL runs. We reasoned that this variability could be explained by the predictor prioritizing salient factors already reliably represented in a random network, thereby impeding feature learning of the less salient factors. To mitigate this tension, we added a mild variance-invariance-covariance (VIC) regularization directly to the encoder’s embeddings (Methods; ***Bardes et al., 2022***; ***Halvagal and Zenke, 2023***). We found that this change led to improved decoding reliability and increased classification accuracy (***Figure 4c***; ***Figure 4—figure Supplement 1***). Consistent with these findings, two-dimensional UMAP projections of the representations revealed that RPL with VIC regularization (RPL^VIC^) led to discernible object category clusters whereas standard RPL and IL did not (***Figure 4d***). The linear projection of representations from an RPL^VIC^ onto the coding directions resembled the ground truth factor representations for most factors, while linear projections of representations from an IL model were mostly collapsed except for the hue (***Figure 4—figure Supplement 2*** and ***Figure 4—figure Supplement 3***). Finally, we confirmed that adding VIC regularization in the IL case did not rescue performance (Supplementary Table S2).

Next, we sought to better understand the comparatively low orientation decoding performance. First, we remind the reader that we use linear probing, in contrast to previous studies that relied on a nonlinear multi-layer network as the decoder (***Garrido et al., 2023***; ***Ghaemi et al., 2024***). Representational untangling requires a common linear coding axis across classes. A necessary, but not sufficient condition of finding such a shared axis is that orientation is reliably encoded for each object individually. Since TLT12 includes objects with diverse shapes and symmetries, we wanted to check to what extent orientation (*θ*_3_) was reliably encoded across object categories. To that end, we trained a set of per-category linear decoders independently (***Figure 4e***). This analysis revealed that RPL networks encoded orientation reliably and markedly better than IL across many, but not all, object categories. Notably, low orientation coding performance correlated with categories with pronounced symmetries, such as earphones, mugs, or smartphones. Similarly, guitars look virtually indistinguishable when viewed from the side (cf. ***Figure 4—figure Supplement 4***). This points to a specific limitation of the linear orientation probing, asserting a 2*π* symmetry, rather than RPL, which prompted us to exclude earphones from the *θ*_3_ decoding analysis. It is remarkable, that despite these difficulties RPL^VIC^ still resulted in partially untangled representations across all remaining categories (cf. ***Figure 4c***; ***Figure 4—figure Supplement 5a, b***). Although the linear projections of RPL^VIC^ representations onto orientation coding directions did not fully resemble the ground truth structure, they showed a structured pattern for *θ*_3_ as well as left/right orientation coding for *θ*_1_, which was not visible in IL projections (***Figure 4—figure Supplement 5c,d,e***).

Finally, we wondered if these representations generalize to novel objects within categories. To this end, we generated an out-of-distribution test set with ten novel random objects within each category. While the resulting representations showed degraded category classification performance for all algorithms, neither RPL nor RPL^VIC^ showed lower performance in decoding dynamic variables (***Figure 4—figure Supplement 6***). Thus, RPL learns untangled representations of dynamic latent variables from more realistic non-deterministic data domains with more factors and changing lighting conditions.

### Representation learning on real-world data

Having established RPL’s effectiveness in extracting latent variables from synthetic datasets, we wondered to what extent these findings would generalize to real-world data. To address this question, we used published video data of behaving mice in an open-field arena (***Luxem et al., 2022***) in which nose position ***r****_N_* and tail root position ***r****_T_* had previously been annotated with DeepLab-Cut (***Mathis et al., 2018***). Based on these annotations, we further estimated animal orientation *θ* and velocity ***v****_T_* (***Figure 5a***; Methods) as latent variables to evaluate the quality of the learned representations.

**Figure 5.**
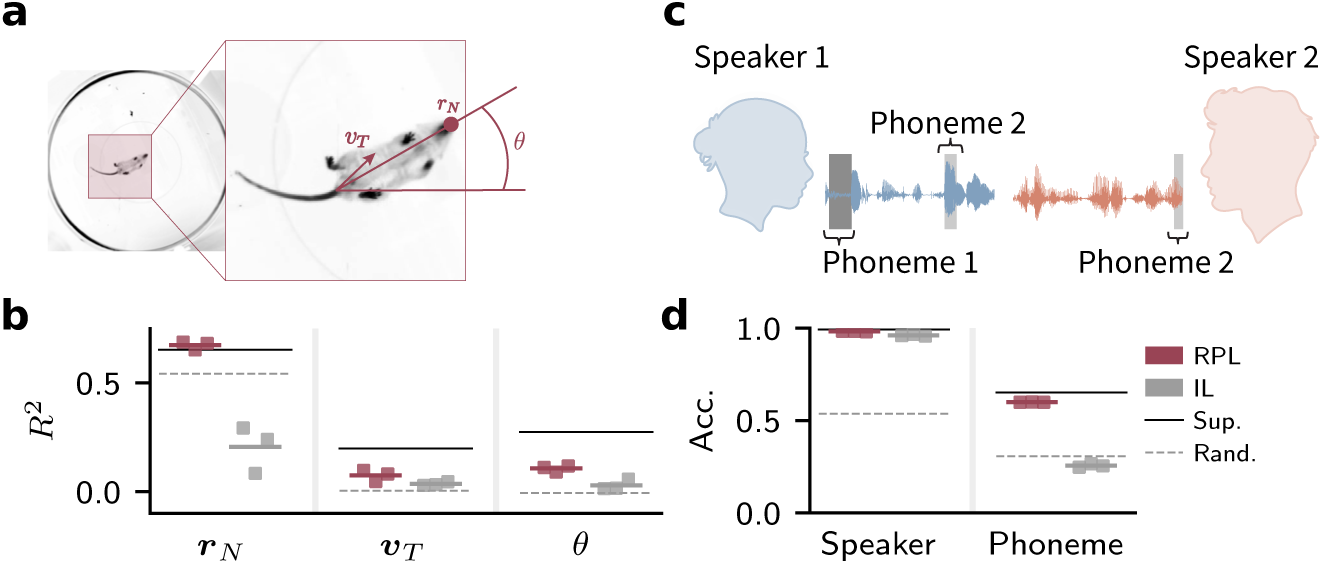
RPL extracts latent variables from real-world data. **(a)** Schematic of real-world video data of behaving mice in an open-field arena (***Luxem et al., 2022***). We used annotated key points to compute position ***r****_N_*, velocity ***v****_T_*, and orientation *θ* as proxies of latent variables for evaluation (Methods). (b) *R*^2^ values of decoded position, velocity, and orientation for RPL and IL from *n* = 3 independently trained networks. Decoding performance for a randomly initialized network (dashed line) and an end-to-end supervised trained network (solid line) shown for reference. (c) Schematic of real-world speech data from the Librispeech corpus (***Panayotov et al., 2015***). We relied on speaker identity and phoneme labels to assess model performance. (d) Linear decoding accuracy of speaker identity and phoneme sequence for RPL and IL for *n* = 3 independently trained models. Decoding from a randomly initialized (dashed line) and an end-to-end supervised trained network (solid line) given for reference. RPL has high decoding accuracy for speaker identity and phonemes. In contrast, IL yields high accuracy only for the speaker identity. ***Figure 5—figure supplement 1***. Latent variable distribution in the mouse data. ***Figure 5—figure supplement 2***. Downstream performance as a function of prediction horizon in the speech dataset.

We then trained RPL networks on the videos and used linear probing for evaluation as before. For comparison, we also trained the same network using the IL objective. We found that decoding from RPL representations yielded higher *R*^2^ values across all variables (***Figure 5b***; Supplementary Table S4), consistent with previous findings, that RPL untangles representations during learning from real-world videos. We observed that velocity was more difficult to decode even when using end-to-end supervised learning, presumably because mice remained stationary in large portions of the videos, leading to a velocity distribution with a peak near zero and a highly imbalanced data (***Figure 5—figure Supplement 1***).

Since the mouse videos were structurally quite similar to our synthetic moving animal videos (cf. ***Figure 2***), we wanted to see whether RPL would yield similarly useful latent representations in a different data domain. To investigate this question, we trained both RPL and IL models on spoken-language recordings from the Librispeech corpus (***Panayotov et al., 2015***). This dataset includes labels for speaker identity and the phonemes of the spoken text (***Figure 5c***). Since phonemes change within and across sentences whilst speaker identity remains unchanged, we expected that RPL would learn representations that allow decoding both, whereas symmetric IL objectives would focus on speaker identity. To check whether this is the case, we evaluated the learned representations by linearly decoding speaker identity and phoneme sequence (Methods). Our results confirmed our expectation. RPL yielded higher accuracy for both speaker identity and phonemes, whereas IL failed to represent phoneme information (***Figure 5d***; Supplementary Table S5). Crucially, the optimal performance in both tasks was achieved with a prediction horizon Δ = 8 corresponding to an average length of a phoneme (∼ 80 ms), suggesting a trade-off between task-relevant and nuisance information as known in contrastive SSL methods (***Tian et al., 2020***; ***Figure 5—figure Supplement 2***). Thus, RPL extracts meaningful latent variables across different real-world data domains.

### RPL learns at different levels of abstraction

In the tasks studied above, the latent-variable dynamics were predominantly smooth in time corresponding to physical object motion or speech. We wondered whether RPL’s specific asymmetric recurrent architecture would enable it to simultaneously learn representations at different levels of abstraction for more diverse stimulus sets and more intricate sequence structures.

To investigate this question, we generated structured sequences of handwritten digits from the MNIST dataset (***Deng, 2012***). Specifically, we generated the sequences using the following grammar. Sequences consisted of distinct digit triplets interspersed by zeros (***Figure 6a***). Only triplets within the clusters of digits {1, 2, 3}, {4, 5, 6}, or {7, 8, 9} were allowed, covering all possible permutations within each cluster. To generate a sequence, we first selected a random cluster, sampled a random permutation within it, and then sampled random instances of the corresponding digits from the MNIST dataset (***Figure 6b***). We continued by sampling the next cluster and triplets according to our defined grammar (Methods). We then fed the resulting image sequences to a network model for training with the RPL and IL objectives.

**Figure 6.**
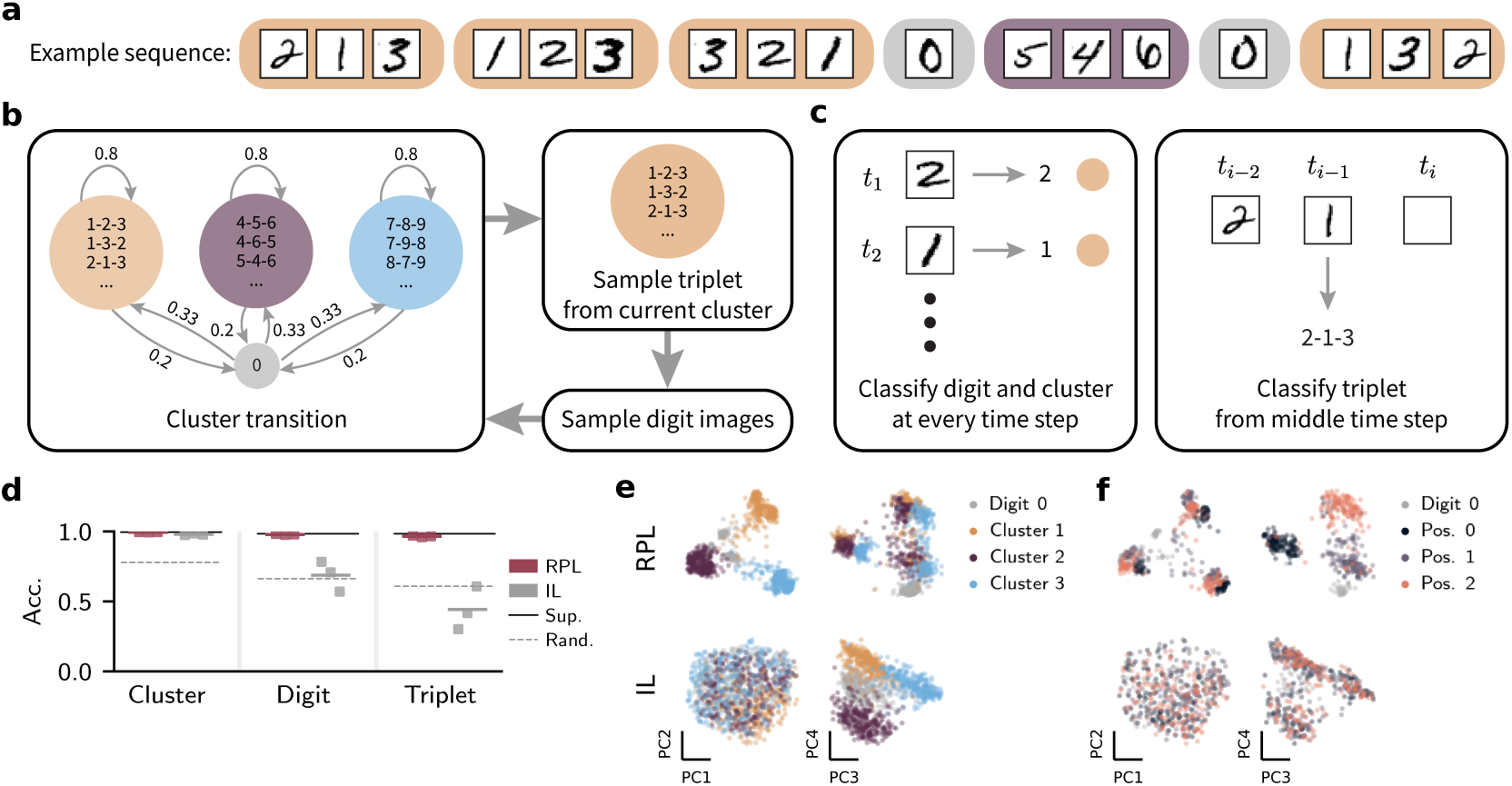
RPL learns at different levels of abstraction. **(a)** An example sequence of digit triplets separated by zeros. **(b)** Schematic of the sequence generation paradigm. We generated triplet sequences containing randomly ordered handwritten digits sampled from one of three digit clusters. Transitions between clusters took place with probability 0.2 accompanied by a “zero”. For any given sequence, each digit instance was randomly chosen from the MNIST dataset. **(c)** Schematic of the tasks used to investigate the emergence of abstraction by asking how well the network representations encode cluster, digit, and triplet identity respectively. Solving these tasks requires both a mapping from images to abstract digit and cluster representations as well as knowledge about the transition structure. To quantify the representational quality, we trained a linear classifier for each task (Methods). **(d)** Linear decoding accuracy of the cluster, digit, and triplet identities for RPL and IL for *n* = 3 independently trained networks. Decoding accuracy for end-to-end supervised learning (solid line) and a randomly initialized network (dashed line) are given for reference. RPL achieves high accuracy on all tasks, whereas IL results in a preference for cluster identity. **(e)** Principal component (PC) projections of the learned representations colored by cluster identity. Both RPL and IL clearly separate the clusters. (f) Same as (e) but colored by temporal position within a triplet. Only RPL yields representations of the abstract temporal transition structure. ***Figure 6—figure supplement 1***. PC projections of the learned representations colored by the different latent variables.

To probe the information encoded in the learned representations, we further trained three sets of linear classifiers to decode digit identity, cluster identity, and triplet identity (***Figure 6c***; Methods). Encoding triplet identity requires a sense of temporal direction and a specific memory of recent digits. In contrast, cluster identity is direction-invariant and does not require distinguishing digits within a cluster. We thus expected RPL, an asymmetric JEPA, to yield high accuracy across all three tasks, in contrast to IL which, due to its symmetric objective, should only perform well at representing cluster identity. This is indeed what we found. RPL-trained network representations solved all three decoding tasks (***Figure 6d***; Supplementary Table S6) with accuracy comparable to that of a fully supervised model.

In contrast, IL resulted in lower-fidelity representations of digit and triplet identity, on par with an untrained random network, while cluster identity was still encoded reliably as expected. Nonetheless, we also found that networks trained with IL could still represent individual digits somewhat faithfully at the encoder level by training a separate set of linear classifiers on the encoder outputs ***z*** (Supplementary Note S2). However, this improved digit-decoding accuracy was still lower than for RPL. This analysis highlights again the importance of asymmetric prediction error computation through the recurrent pathway for learning time-resolved and abstract representations, even when these representations themselves do not depend on temporal context, as is the case for individual digit stimuli.

To gain further insight into how the temporal context is encoded in the trained networks, we visualized the principal components (PCs) of the learned representations (***Figure 6e,f***; ***Figure 6—figure Supplement 1***). While cluster identity was clearly separated by both methods, e.g., along the first two PCs for RPL, and third and fourth PCs for IL (***Figure 6e***), it was evident that RPL resulted in more finely structured representations. For instance, RPL representations encoded the temporal position within a triplet in the third and fourth PCs, essentially creating an internal clock that tracks sequence progression (***Figure 6f***). Furthermore, PCs five through eight displayed a rich, though less interpretable, organization (***Figure 6—figure Supplement 1***), presumably encoding the digit identity. In contrast, IL lacked such a fine-grained organization beyond clearly separating cluster identity, lending further support to our hypothesis that RPL’s asymmetric architecture is essential for learning abstract sequence structure that requires a sense of direction, whereas IL’s symmetric setup is limited to representing invariant features. Thus, RPL’s asymmetric recurrent architecture is necessary to simultaneously learn complex representations at different abstraction levels from abstract sequence stimuli.

### RPL learns successor-like representations comparable to V1 in humans

We next wondered to what extent RPL network models predicted *in vivo* data. To that end, we first focused on a specific class of predictive representations: successor representations. In their original, reinforcement-learning formulation, successor representations describe how a neuronal population encodes a *predictive* representation of future states, such that each state representation reflects the expected, policy-dependent, discounted future occupancy of other states (***Dayan, 1993***). RPL training involves no reward, value, or policy: it is a purely predictive, self-supervised objective operating on passively observed sequences. We therefore do not ask whether RPL learns successor representations in the strict reinforcement-learning sense; rather, following ***Ekman et al. (2023)***, who coined the term *successor-like* representations to describe an analogous signature in a similarly policy-free, passive-viewing setting, we use it here to refer specifically to the characteristic *asymmetric, temporally-ordered* pattern of predictive activation across sequence positions, and not to the full discounted, policy-dependent quantity defined by Dayan (1993). ***Ekman et al. (2023)*** described such *successor-like* representations in fMRI recordings from the primary visual cortex of humans (***Ekman et al., 2023***). During these experiments, humans were exposed to a sequence of four non-overlapping items (A-B-C-D) across multiple trials. After an initial learning phase, the participants were exposed to either the entire sequence or only one of the four sequence items, referred to as partial sequences (***Figure 7a***). The data from V1 indicated activation at the successor but not the predecessor locations upon partial sequence exposures (***Figure 7b***).

**Figure 7.**
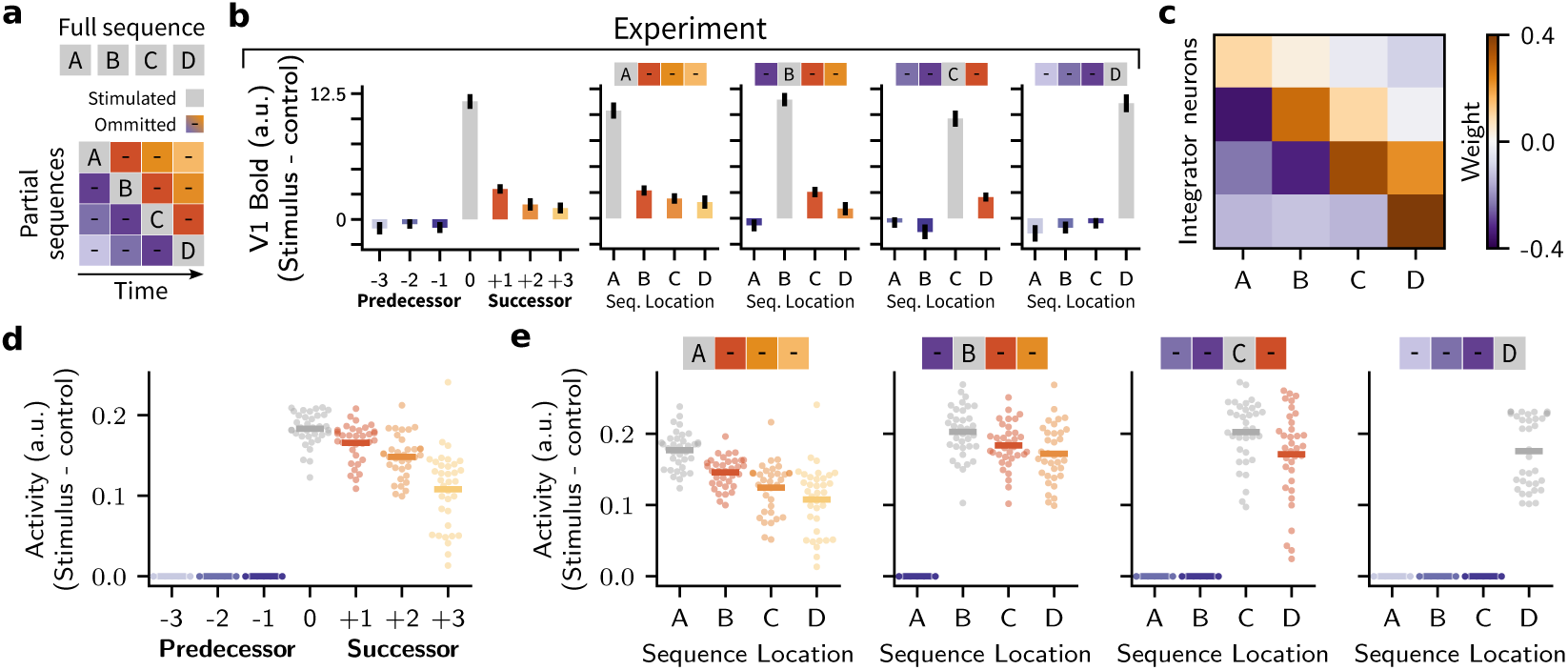
RPL learns successor-like representations comparable to human V1. **(a)** Schematic of the full and partial sequences used in the experiment. (b) Relative BOLD signal responses reproduced in human V1 (***Ekman et al., 2023***). Relative stimulus response to partial sequence stimuli (left). Individual responses to partial sequences (right). Partial sequence stimulation caused brain activity encoding successor locations but not predecessor locations. Error bars represent the standard error of the mean. Data extracted and reproduced from ***Ekman et al. (2023)***. (c) Relative neuronal selectivity matrix extracted from the RPL model representations (*n* = 35 models). The negative values below the diagonal indicate inhibition of activity at predecessor locations. Positive values above the diagonal indicate that the successors’ representations are driven by the stimulus. (d) Average neural activity for all partial sequences at non-stimulated and stimulated locations after removing the baseline activity as in (b). Each dot is a trained model and the solid lines represent the average activity over all models. The plot indicates positive neural activity at stimulated and successor locations, and no activity at predecessor locations. (e) Neural activity at all sequence locations in response to each partial sequence. These plots indicate the decaying positive response at successor locations for individual partial sequences, hinting at successor-like representations in response to each partial sequence.

We wanted to know whether RPL networks exhibit similar activation patterns at successor locations after being exposed to comparable sequential stimuli. To investigate this question, we trained minimal RPL circuit models (*n* = 35; Methods) on an (A-B-C-D) sequence to mirror the experimental protocol. After sorting the neurons in the integrator based on their selectivity, we visualized the average weight matrix associated with the neural selectivity to each sequence item (***Figure 7c***). This matrix showed the largest entries on the main diagonal, positive values on the two upper off-diagonals, and negative values below the main diagonal, a clear signature of successor-like representations. Again, without reward or policy, this asymmetric structure is equivalent to the network’s learned representation of the sequence’s transition structure, i.e., a primitive, policy-free predictive model of “what comes next,” rather than a full successor representation, which is why we refer to it as *successor-like* for consistency with ***Ekman et al. (2023)*** and to facilitate direct comparison with their V1 data.

We then computed the average neural activity at the stimulated, predecessor, and successor locations in response to partial sequences, after removing baseline neural activity from all neural responses. We observed that, as in the fMRI recordings (***Ekman et al., 2023***), the stimulated location showed the highest activity, while activity at successor locations followed a decaying pattern. In contrast, there was no activity at the predecessor locations consistent with the experimental data (***Figure 7b,d***). Similarly, we plotted neural activity in response to each partial sequence to see whether this pattern also existed in selective neurons for non-stimulated sequence items at successor locations. We observed a consistent pattern of activity at stimulus and successor but not predecessor locations (***Figure 7e***). Thus, RPL entails *successor-like* representations as observed in V1 of humans experiencing repeating sequence stimuli.

### RPL learns abstract sequence representations similar to macaque PFC

Having observed that RPL learns predictive representations consistent with those in a primary sensory cortex, we asked whether our model could also capture more abstract representations ascribed to higher-order cortical areas, such as the PFC. A widely used experimental setup to investigate abstract sequence learning is the local-global oddball paradigm (***Figure 8a***). Here, an animal is exposed to trials or sequence presentations consisting of four stimuli one after the other, either repeating the same stimulus (xxxx or xx for short) or an oddball in the last slot (xxxY or xY for short), constituting a local deviant. Repeating trials are organized into blocks, in which the animal is conditioned to expect a particular sequence type, e.g., xx or xY, as the standard by dint of frequent presentation at the beginning of the block. Later trials in the block will be either global standards or infrequent global deviants, depending on whether they match the context set at the start of the block. For example, in a block where xY is the standard, an xx trial denoted (xx xY) constitutes a global deviant but not a local one.

**Figure 8.**
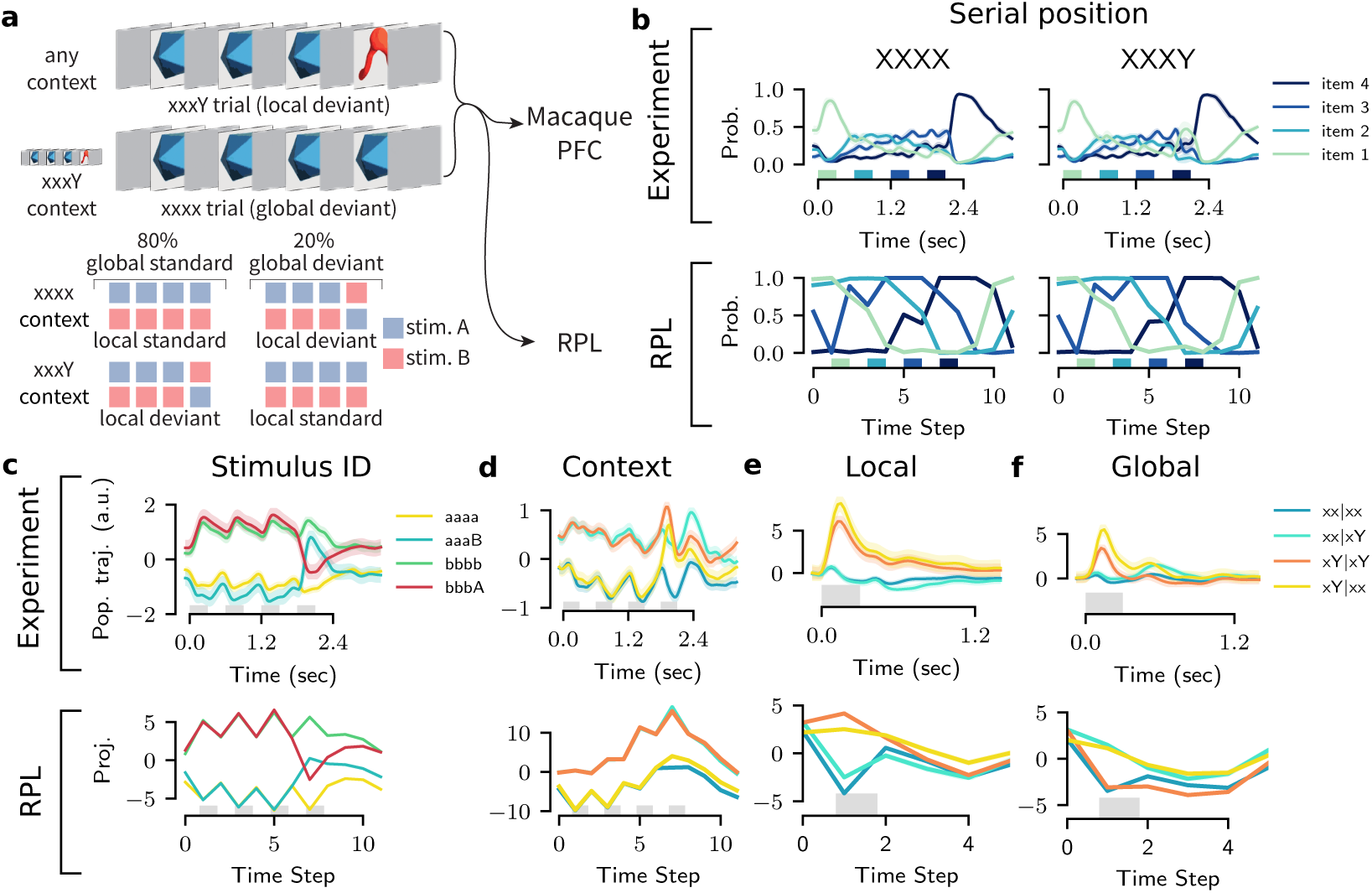
RPL learns abstract sequence representations similar to macaque PFC. **(a)** Schematic of the local-global oddball paradigm used by ***Bellet et al. (2024)***. Example trials consisting of four sequential stimuli show an “xxxY” and an “xxxx” trial in the context of frequent “xxxY” presentations (top). A given trial can be a local standard or deviant depending on the local sequence structure of the four stimuli and a global standard or deviant depending on the context set for a block of several trials (bottom). Figure adapted from ***Bellet et al. (2024)***. (b) Comparison of temporal serial position representation in experiment (top) and the model (bottom). Plots show best linearly decoded probabilities of the four serial positions (items 1 through 4) averaged over all xxxx and xxxY trials. RPL learns representations that encode serial position more robustly than the macaque PFC. The experimental data corresponds to Monkey A and has been reproduced from publicly available data and code. (c) Comparison of stimulus identity representation in the experiment (top) and the model (bottom). Plots show projections of neural activity or RPL representations on directions that best separate pairs of input stimuli (Item a vs b) that comprise x and Y, averaged across trials (Methods). RPL representations encode stimulus identity similar to the experiment. (d) Same as (c) but for representation of the global context set by the frequent trial. (e) Same as (c) but for the presence or absence of a local deviant at the fourth sequence position. (f) Same as (c) but for the presence or absence of a global deviant, decoded from the fourth sequence position.

One such experiment was performed by ***Bellet et al. (2024)*** who recorded neurons from the macaque PFC using multielectrode Utah arrays while exposing them to visual local-global oddball sequences. They found that populations of neurons in the PFC spontaneously learned to encode several abstract sequence properties, such as their serial position within a trial of four stimuli, stimulus identity, global context, and the presence of local and global deviants.

Since we had already observed that RPL is capable of abstract sequence learning (cf. ***Figure 6***), we hypothesized that RPL should also be able to learn the sequence properties of local-global oddball sequences. To test this hypothesis, we trained an RPL circuit model on the identical visual stimulus sequences that ***Bellet et al. (2024)*** presented to the macaques, closely mirroring their experimental protocol. Then, following their analysis (Methods), we extracted neuronal population vectors by training linear readouts to decode serial position, stimulus identity, global context, and local/global deviance from RPL representations ***c***.

We found that the serial position of an item in a sequence and stimulus identity could be decoded reliably (***Figure 8b,c***) as in the experiments, albeit without the experimentally observed accentuation of the first and the last item, which are presumably due to primacy and recency effects in the experiment (***Bellet et al., 2024***). To test whether the RPL network had learned an internal model of the sequence structure, we decoded the global context variable from the internal representations. As observed in the authors’ original analysis, we identified neural subspaces in our model that coded for the global context inferred from previous sequence observations (***Figure 8d***). Finally, we assessed our model’s responses to local and global deviants. We found qualitatively similar separation of population responses corresponding to the presence of local deviance in both predicted (xY|xY) as well as unpredicted (xY|xx) local deviant stimuli (***Figure 8e***). Similarly, the model showed an explicit encoding of the presence of global deviance (xx|xY and xY|xx; ***Figure 8f***) in its population responses, as in the experiment. Although our model did not exhibit the onset and primacy effects observed in the data, presumably because it lacked neuronal adaptation mechanisms, we found that the representations learned via RPL qualitatively matched the relative encoding in PFC. Thus, RPL learns abstract sequence representations whose relative encoding structure is consistent with PFC activity.

### Untangling in a cortical model with local prediction error circuits

So far, we have used deep encoder networks, computing prediction errors at the top-level output, using backpropagation of error (backprop) to update all weights. However, experimental evidence points to local mismatch computation across the hierarchy as early as primary sensory cortex (***Keller and Mrsic-Flogel, 2018***). Moreover, how the brain approximates backprop to solve the credit assignment problem remains an open question (***Lillicrap et al., 2020***). Hence, we wondered whether RPL remains effective without end-to-end optimization and to what extent.

To investigate this question, we built a hierarchical RPL (h-RPL) model in which we implemented different cortical areas as local RPL circuits that learn independently. Functionally, each circuit consists of a shallow encoder, an integrator, and a predictor putatively implemented by distinct cell types. One possible mapping of the model onto cortical anatomy is to view the encoder as being implemented through lateral connectivity between adjacent areas, whereas vertical inter-laminar connectivity implements the integrator and predictor, resulting in a circuit that produces local prediction errors (***Figure 9a***) reminiscent of mismatch responses observed in superficial cortical layers (***Keller and Mrsic-Flogel, 2018***).

**Figure 9.**
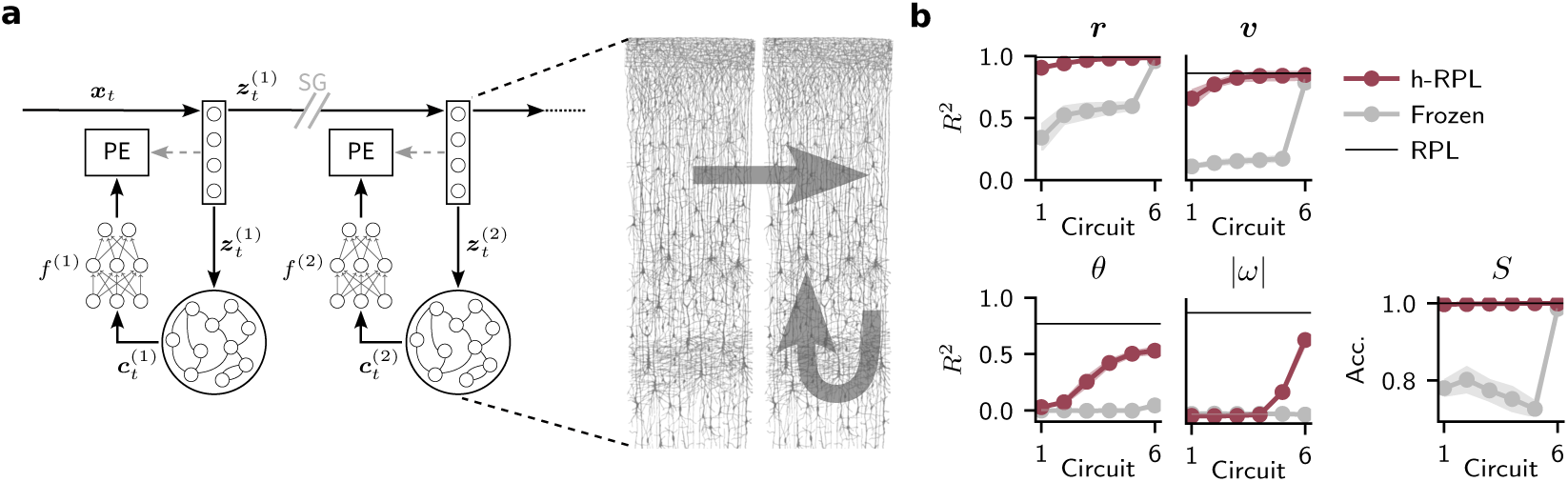
Untangling in a cortical model with local prediction error circuits. **(a)** Schematic of the network model consisting of six identical RPL circuits in a hierarchy, each containing a shallow encoder, an integrator and a predictor network. Each circuit computes its own prediction error while Stop Grad (SG) operations prevented gradients from flowing between the individual circuits. In this h-RPL model horizontal intra-laminar connectivity implements the encoder network. In contrast, vertical connectivity accounts for the integrator and predictor circuitry. Cortical anatomy drawing (right) adapted from Cajal (***y Cajal, 1899***). **(b)** Decoding *R*^2^ values and accuracy of the latent variables (cf. ***Figure 2b,c***) for each RPL circuit in h-RPL (red) and a variant in which only the last circuit in the hierarchy was trained (gray). Shading corresponds to one standard deviation over *n* = 3 independent models. The solid black line denotes the performance of RPL with a deep encoder (cf. ***Figure 2j***). The h-RPL model progressively learns latent variable representations comparable to end-to-end trained RPL. ***Figure 9—figure supplement 1***. Straightness of learned representations increases over the hierarchy and over learning. ***Figure 9—figure supplement 2***. Autocorrelations of learned representations increase over the hierarchy and over learning.

We trained this h-RPL model on synthetic moving animal data (cf. ***Figure 2***). For simplicity, we trained every circuit using backprop, but we prevented gradients from flowing between circuits across the hierarchy (Methods). Doing so confines credit assignment to individual circuits with a few synaptic transmission steps, a setting in which biologically plausible backprop approximations are effective (***Lillicrap et al., 2016***; ***Nøkland, 2016***; ***Whittington and Bogacz, 2017***; ***Sacramento et al., 2018***). In consequence, each circuit *l* learned independently in this hierarchical network model.

To assess the quality of the learned representations, we trained linear decoders for each circuit and extracted the latent variables as before (***Figure 9b***). This analysis revealed that the linear decoding accuracies and *R*^2^ values for the dynamic variables increased with depth along the hierarchy, plateauing at performance levels comparable to those of the end-to-end trained RPL setup. To check whether this linear decodability of the evolving latent variables also implied “perceptual straightening,” a phenomenon characterized by the progressive transformation of curved stimulus trajectories into more linearly evolving representations along the visual cortical hierarchy (***Hénaff et al., 2019***, ***2021***), we further quantified straightness along the model’s hierarchy. We found, that straightness indeed increased along the hierarchy and was due to learning (***Figure 9—figure Supplement 1***).

One potential caveat with this analysis is that the hierarchical encoders jointly constitute a deep convolutional network with increasing receptive field sizes. To check that the observed effects were not merely due to the progressively larger receptive fields, we trained another instance of the same hierarchical model in which we kept the weights in all but the last circuit of the hierarchy frozen. This resulted in an overall reduction in linear decodability across the hierarchy (***Figure 9b***; ***Figure 9—figure Supplement 1***). While the drop in *R*^2^ for position, velocity, and object category could be almost entirely compensated for by the optimized last circuit, this was not the case for orientation and angular speed. Additionally, we observed a steady increase in the autocorrelation time of the representations over the hierarchy for h-RPL but not in the frozen counterpart (***Figure 9—figure Supplement 2***). These findings are consistent with the idea that better decodability across circuits is not merely a result of increasing receptive field size, but rather the progressive untangling of representations across the hierarchy. Thus, RPL is a powerful learning algorithm for finding dynamic latent variables even in a hierarchical setting, which resonates with the core ideas of predictive processing and the experimentally observed mismatch neurons in primary sensory cortices.

## Discussion

We introduced recurrent predictive learning (RPL), a self-supervised learning model that learns untangled representations of objects and their dynamics directly from sequential observations alone, without requiring labels, an implausible decoder, or a reconstruction objective. Formally, RPL constitutes a JEPA that extracts both invariant and “equivariant” representations across multiple levels of abstraction from synthetic and real-world data. Beyond representation learning, RPL also acquires an internal world model that can simulate plausible future states purely based on its internal representations. Crucially, the learned internal representations resemble neuronal activity patterns reported in macaque PFC and exhibit successor-like signatures comparable to those observed in human primary visual cortex.

In this article, we use the term *equivariant representation* loosely, in a broader sense, to denote a representation that jointly encodes latent variables and their temporal dynamics. This framing applies naturally to continuous latent variables, such as an object’s position or orientation, where a well-defined rate of change exists. For discrete or abstract sequences, such as the non-overlapping symbolic sequences used in ***Figure 6***, latent states change categorically between discrete items rather than continuously. Here we mean the underlying predictive relationship between successive states rather than a rate in the strict sense. This joint encoding is functionally motivated: predicting future latent states requires not only *what* is currently present (captured by invariant content) but also *how* it is changing, since the two together determine the system’s trajectory forward in time. It should be noted that this notion of equivariance differs from the stricter standard usage in machine learning, where equivariance refers to a *memoryless* mapping whose output transforms predictably under a given transformation of its input (e.g., *f*(*gx*) = *g*^′^*f*(*x*)), without any requirement that the transformation’s rate or dynamics themselves be explicitly represented. In RPL, by contrast, the recurrent integrator accumulates evidence across time, so that its state represents not only the current value of a latent variable but also how quickly and in which direction it is changing, which is precisely the information the circuit needs to support prediction.

Learning in RPL is driven by internal prediction errors generated within a specific circuit motif: an encoder with little working memory feeds into an integrator-predictor circuit that integrates temporal information and learns to anticipate future encoder embeddings. This motif appears functionally general, enabling hierarchical networks with local predictor circuits to progressively uncover latent variables. While invariant representations can be learned by local learning rules in conjunction with appropriate decorrelation mechanisms (***Halvagal and Zenke, 2023***), we showed in this work that learning representations of dynamic variables requires a circuit with distinct functional elements for encoding, integration, and prediction. We expect analogous functional motifs to exist in brain areas that rely on self-supervised learning, implemented through dedicated cell types and connectivity patterns that jointly shape plasticity. In this way, our work provides a conceptual blueprint for interpreting neuronal circuits, while making concrete, testable predictions.

### Relation to previous modeling work

The notion that the brain builds and progressively updates an internal model of the world through comparison with observations is a central tenet of systems neuroscience (***Rao and Ballard, 1999***; ***Bastos et al., 2012***; ***Keller and Mrsic-Flogel, 2018***; ***Recanatesi et al., 2021***; ***Hawkins, 2021***; ***Illing et al., 2021***; ***Brunec and Momennejad, 2022***; ***Price et al., 2023***; ***Tang et al., 2023***; ***Levenstein et al., 2024***; ***Gillon et al., 2024***; ***Richter et al., 2024***; ***Han and Helmchen, 2024***). Classical predictive coding models based on this idea assume an underlying generative model that *reconstructs* sensory input (***Rao and Ballard, 1999***; ***Lotter et al., 2020***; ***Hertäg and Sprekeler, 2020***; ***Mikulasch et al., 2023***; ***Boeshertz and Clopath, 2025***). The generative model is learned and updated by computing prediction errors in the input space. However, classical predictive coding is under increasing pressure due to its difficulty in explaining recent experimental data (***Solomon et al., 2021***; ***Richter et al., 2024***; ***Westerberg and Roelfsema, 2025***); moreover, it yields poor representations for discriminating between invariant concepts (***Oliviers et al., 2025***). Modern variants of predictive coding alleviate this problem by combining a supervised loss function with the classic unsupervised reconstruction loss (***Oliviers et al., 2025***). Still, doing so requires labels. These difficulties might stem from its core assumption of an underlying generative model. Indeed, recent studies argue that reconstruction, a core component of generative models, is inefficient for learning high-level abstract representations because it tends to focus on behaviorally irrelevant low-level stimulus details (***Balestriero and LeCun, 2024***; ***Van Assel et al., 2025***).

In contrast, JEPAs like RPL depart altogether from the idea of an underlying generative model. Nevertheless, RPL, and h-RPL in particular, share some essential elements with classical hierarchical predictive coding models. In all cases, learning is driven by prediction errors that are computed locally and used to update model parameters and internal representations, a mechanism consistent with the existence of prediction-error-like neurons in sensory cortex (***Zmarz and Keller, 2016***; ***Keller and Mrsic-Flogel, 2018***; ***Gillon et al., 2024***). In this respect, h-RPL aligns with established predictive coding models in which cortical circuits continuously generate and refine predictions about incoming signals (***Rao and Ballard, 1999***; ***Lotter et al., 2020***; ***Li et al., 2025***).

However, a central distinction lies in what is communicated between the circuit elements. Classical predictive coding architectures rely on the explicit propagation of prediction errors across hierarchical levels, typically assuming a functional segregation between representation neurons and error neurons. In contrast, the JEPA-like organization underlying h-RPL does not require prediction errors to be transmitted beyond the local circuit in which they are computed. Instead, separate network structures embed related signals and learn to predict each other’s latent states via locally computed errors. As a result, inter-areal communication is dominated by representations rather than error signals, with learning occurring through the mutual alignment of internal embeddings. This shift fundamentally changes the representational objective of the network, emphasizing abstract, predictive latent variables over the reconstruction of sensory input. Finally, we should note that this shift is not mutually exclusive with modern forms of *discriminative* predictive coding, which, in conjunction with a predictive self-supervised loss, could support representation learning by approximating backprop (***Whittington and Bogacz, 2017***; ***Oliviers et al., 2025***).

Based on these arguments, this article advocates for a revised predictive processing framework, one that avoids explicit generative models, meaning that it neither relies on reconstruction in the input space nor supervised objectives, and instead relies on SSL via prediction in the representation space. Recent studies have taken steps in this direction by developing reconstruction-free predictive SSL models that compute predictions directly in the embedding or representation space. For example, ***Ororbia et al. (2025)*** proposed a model that relies on spatial prediction across visual scene embeddings. Other models rely on contrastive learning (***Illing et al., 2021***; ***Delrocq et al., 2024***) or temporal prediction (***Halvagal and Zenke, 2023***) across sequences of augmented images reminiscent of slow feature analysis (***Wiskott and Sejnowski, 2002***). These studies have provided valuable insights into learning invariant representations without supervision. However, they primarily focused on category-level invariance induced by hand-crafted augmentations and did not explicitly address how biological agents may simultaneously acquire equivariant representations that preserve behaviorally relevant information, such as motion direction or relative spatial relationships.

Several recent studies have proposed related models of the visual system that operate on sequential observations (***Nejad et al., 2025***; ***Niu et al., 2024***; ***Bakhtiari et al., 2021***). For instance, ***Bakhtiari et al. (2021)*** suggest that functional specialization can arise in a two-pathway network trained with a single predictive contrastive objective, though the extent to which dynamic variables are explicitly represented remains an open question. ***Niu et al. (2024)***, in turn, propose a temporal “straightening” objective as an alternative to invariance for learning equivariant features – a complementary perspective to ours. However, since the resulting equivariant representations are evaluated using nonlinear decoders, it remains unclear to what extent the underlying latent variables are untangled and how the straightening objective could be implemented across cortical hierarchies.

Related to our work, ***Nejad et al. (2025)*** introduced a temporal predictive SSL model with a putative mapping onto neocortical layers. In this model, prediction occurs in the latent space spanned by Layer 5 (L5). Specifically, Layers 2/3 (L2/3) predict the next L5 representation. To avoid representational collapse, L5 is either trained like an autoencoder with an additional reconstruction objective, or by adding a variance-maximization criterion. The resulting model learns to predict orientation changes while capturing several experimental observations. In contrast to RPL, the study used shallow encoder networks and utilized sequential Gabor patches as sensory input, with the rotation direction provided as top-down input. Similarly, ***Asabuki et al. (2025)*** investigated predictive sequence learning in a local recurrent circuit model, emphasizing biologically plausible learning rules. While these studies make notable contributions toward understanding how brains may implement predictive learning, they focused on comparatively simple network architectures and elementary learning tasks. They did not address the emergence of rich, abstract representations from complex sensory streams.

While the above studies similarly argue for a set of core principles that should underpin a biologically grounded theory of representation learning, i.e., one that does neither rely on reconstruction nor supervision, the same principles naturally align with the class of SSL models developed in machine learning under the umbrella of JEPAs (***Noroozi and Favaro, 2016***; ***Oord et al., 2018***; ***Grill et al., 2020***; ***Zbontar et al., 2021***). JEPAs operationalize the idea that internal representations can be learned by making different views or temporal segments of sensory experience predict one another, thereby preserving the core intuition of predictive processing while avoiding many of the constraints imposed by explicit generative models. Crucially, RPL is a JEPA, which means it neither relies on a reconstruction objective nor on negative samples, and by avoiding the issues associated with generative models, it learns useful abstract representations from a stream of sensory experience, all the while remaining consistent with the core idea of predictive processing and neural circuit principles.

Although a plethora of different JEPAs have demonstrated their effectiveness in machine learning (***Balestriero and LeCun, 2024***; ***Van Assel et al., 2025***; ***Halvagal and Zenke, 2023***; ***Niu et al., 2024***; ***Grill et al., 2020***; ***Chen and He, 2021***; ***Assran et al., 2023***; ***Ghaemi et al., 2024***; ***Bardes et al., 2024***; ***Assran et al., 2025***), not all of them are biologically plausible. For instance, existing studies made strong assumptions about the allowed transformations between consecutive stimuli (***Ghaemi et al., 2024***) or were not applied to real-world problems and lacked a circuit-level interpretation (***Richthofer and Wiskott, 2015***). In part, this lack stems from current JEPAs typically relying on biologically implausible transformer networks, predicting randomly masked parts of the stimulus, and optimizing a single global prediction error during training, all of which are difficult to reconcile with neural anatomy. Finally, the majority of research has focused on invariance learning, in which representational quality is assessed primarily through object classification tasks, e.g., via linear decoding or probing. Such quantification of model performance at the level of internal representations is a welcome and necessary departure from evaluating representational quality solely on a model’s ability to reconstruct stimuli at the pixel level, as commonly done for classical predictive coding models (***Vladimirskiy et al., 2015***; ***Lotter et al., 2020***). However, focusing solely on invariance learning seems like an oversimplification. An increasing number of machine learning studies are acknowledging the importance of equivariant representations (***Devillers and Lefort, 2022***; ***Park et al., 2022***; ***Garrido et al., 2023***; ***Wang et al., 2024***; ***Ghaemi et al., 2024***; ***Niu et al., 2024***; ***Drozdov et al., 2024***), albeit with the limitations mentioned above and the assumption of direct access to the underlying transformations. A notable departure from these approaches is ***Drozdov et al. (2024)***, who propose a temporal JEPA with additional variance-covariance regularization, framed within an energy-based learning framework. This work represents an important step away from transformer-based masked-representation-prediction JEPAs; further evidence would nonetheless help establish the extent to which the learned representations are untangled and equivariant.

RPL avoids several implausible aspects of the above studies. On the one hand, RPL is a temporal JEPA based on recurrent neural networks rather than transformers. On the other hand, it does not rely on masking as commonly used in self-supervised video models (***Bardes et al., 2024***; ***Assran et al., 2025***). ***Velarde and Parra (2024)*** studied a related recurrent JEPA while focusing on biologically motivated learning rules that avoid backprop through time. While similar to our approach, their setup closely resembles our CL objective, which proved less effective in our study. Finally, h-RPL is, to the best of our knowledge, the first implementation of a hierarchical JEPA that optimizes multi-level prediction errors as initially proposed by ***LeCun (2022)***. Functionally, RPL learns both invariant and equivariant representations consistent with the idea of active filtering (***Histed, 2025***), as well as a latent generative model that enables it to simulate plausible dynamics in the latent space when queried with its own predictions. Interestingly, the necessity of separating the feedforward prediction target from the integrator-predictor branch directly stems from this reliance on a plausible recurrent neural network architecture, which opens the door to further investigation of the putative links between our model and the cortical microcircuit.

Lastly, avoiding representational collapse is a challenge in SSL models and puts additional pressure on the algorithms that the brain can implement. JEPAs are particularly interesting biologically plausible candidates as they can prevent representational collapse through an asymmetric vector-based prediction target (Methods; ***Chen and He, 2021***; ***Tian et al., 2021***; ***Srinath Halvagal et al., 2023***). Whether and how biological circuits use such vector-based targets for learning remains an open question, but several theoretical proposals exist of how they could (***Rossbroich and Zenke, 2025***; ***Laborieux and Zenke, 2022***; ***Lillicrap et al., 2020***; ***Song et al., 2024***; ***Sacramento et al., 2018***). While many JEPAs rely on secondary mechanisms to avoid trivial solutions or collapse (***Grill et al., 2020***; ***Assran et al., 2023***; ***Bardes et al., 2024***; ***Assran et al., 2025***; ***Drozdov et al., 2024***; ***Balestriero and LeCun, 2025***), RPL did not require such auxiliary mechanisms in the situations we tested. One notable exception were our tests on TLT12, where we observed that a weak regularization on the embeddings could further stabilize learning (cf. ***Figure 4c***). Other than that, RPL follows conventional JEPA design principles, whereby the predictor network learns at a higher learning rate (***Chen and He, 2021***). Recent theoretical work (***Mikulasch and Zenke, 2026***) linked the predictor output to the network’s conditional entropy estimate, which must not collapse, and that the learning rate difference is key for keeping this entropy estimate from becoming stale. Finally, SSL algorithms entail a trade-off between perceptually relevant and nuisance information in the predicted and target embeddings (***Tian et al., 2020***). We empirically showed that a similar trade-off, stemming from the prediction time horizon Δ, exists in RPL (cf. ***Figure 5—figure Supplement 2***). Collectively, these mechanisms seem prevent RPL from learning trivial solutions while remaining simple and implementable in biological networks.

In the hierarchical setting, one may wonder why circuits higher in the hierarchy learn increasingly abstract features. We believe two mechanisms are at play here: First, the hierarchical encoder is a stacked convolutional network, meaning that receptive field sizes grow with depth. This architectural inductive bias naturally encourages successive circuits to operate on increasingly spatially integrated features, creating a structural pressure toward learning complementary rather than redundant representations. Second, the growing expressivity of the network with depth means that higher circuits have access to richer, more abstract inputs from which they can extract higher-level latent structure that is not already captured by lower circuits. Together these factors: the local collapse-preventing mechanism and the depth-dependent growth in receptive field size and network expressivity presumably explain why h-RPL builds an increasingly refined and non-redundant representational hierarchy.

### Relation to experimental observations and circuit structure

Our work reveals several points of convergence between RPL and experimental observations of cortical computation, spanning both early sensory and higher-order areas. At a functional level, RPL develops representations that are explicitly optimized for prediction over time, rather than reconstructing sensory input. In particular, RPL learns successor-like sequence representations, in which current stimuli evoke activity at future but not preceding locations, closely paralleling successor representations recently reported in human primary visual cortex (***Ekman et al., 2023***) and broadly consistent with sequential activity recall observed in mouse V1 (***Xu et al., 2012***). Beyond early sensory areas, RPL also acquires abstract sequence representations that resemble those observed in macaque PFC during local–global oddball paradigms (***Bellet et al., 2024***). These representations encode relational and temporal structure rather than sensory detail, consistent with the idea that higher cortical areas operate on increasingly abstract latent variables.

What makes these correspondences particularly noteworthy is that we fit no model parameters to the neural recordings in either case. Rather, the representational signatures emerge naturally when RPL is exposed to the same temporal sequences used in the respective experiments. We selected both studies for comparison because they involve passive observation of temporal sequences without an explicit task, closely matching the self-supervised setting in which RPL is trained. That a single architecture with a simple predictive learning rule reproduces key representational signatures across two anatomically and functionally distinct brain areas, spanning early sensory cortex and higher-order association areas, provides a strong sufficiency argument for RPL as a candidate theory of cortical representation learning.

Moreover, our results align with experimental and theoretical work suggesting that cortical representations are shaped by predictive objectives (***Palmer et al., 2015***; ***Singer et al., 2018***; ***Taylor et al., 2024***) and become progressively more invariant and abstract over the cortical hierarchy (***Piasini et al., 2021***). In h-RPL, we observed a similar systematic increase in temporal stability across circuits (***Figure 9—figure Supplement 2***), mirroring timescale hierarchies observed across cortical areas (***Murray et al., 2014***). Finally, h-RPL offers a mechanistic account of perceptual straightening phenomena in visual cortex (***Hénaff et al., 2019***, ***2021***), which refers to the progressive reorganization of sensory representations such that temporally structured or correlated stimuli come to be represented as more stable, linearized trajectories in neural state space, improving predictability and downstream readout. Our work suggests that such effects may arise naturally from representation-space predictive learning rather than from explicit sensory reconstruction.

In relation to cortical anatomy, the central contribution of our model is not a one-to-one mapping between algorithmic components and specific cortical cell types or layers, but the proposal that representation learning of invariant and equivariant representations requires a circuit with specific elements: encoder, temporal integrator, predictor, and prediction-error computation, all implemented by different circuit elements and, ultimately, different neuronal populations. This separation of function is a defining feature of RPL and of JEPAs more generally. Importantly, it does not require that these components be realized in a single, unique, or canonical circuit layout.

In the minimal model studied here, these elements are arranged in a feedforward–recurrent motif without explicit top-down connections or recurrent interactions between circuit modules. This abstraction is an oversimplification of cortical reality, which was chosen deliberately to isolate the core circuit elements for predictive learning using internal representations, not to provide a complete account of cortical anatomy. The cortex contains dense recurrent connectivity within and across layers, extensive lateral interactions, and bidirectional long-range projections, any of which could support alternative implementations of the same underlying computation. As a result, the precise correspondence between model components and cortical microcircuits remains inherently ambiguous.

That said, an appealing idea that is broadly consistent with our h-RPL model is that the encoder could be implemented through lateral intra-layer connectivity, whereas the circuits required for generating predictions and prediction errors follow a vertical columnar organization. In this context, L2/3 neurons, which have been associated with prediction-error-like signals, are a prime candidate for computing prediction errors. Still, how the integrator and predictor map onto the cortical circuitry is less clear. Even if these elements did follow a vertical organization, they could well be intermingled without clear laminar division. Finally, while our RPL implementation assumes a single encoder, the JEPA framework naturally admits alternative configurations with distinct encoder networks (***Grill et al., 2020***), which could reside in the superficial and deep layers of the cortex. This ambiguity underscores that the framework specifies which computations must be present, rather than where they must reside, and highlights the need for future experimental work to disambiguate among these possible biological realizations.

Another notable feature of h-RPL is that prediction error acts as a learning signal rather than a representation that is being feedforward communicated. While this property is at odds with classical predictive coding (***Rao and Ballard, 1999***), recent neurophysiological evidence suggests that residual computation may not be the primary coding principle in cortex. For instance, ***Furutachi et al. (2024)*** observed selective amplification of responses in mouse primary visual cortex, leading to a revised framework in which sensory representations remain the main mode of feedforward communication, modulated by prediction error signals (***Furutachi and Hofer, 2026***). h-RPL is compatible with such representational feedforward communication. Recently, ***Vasilevskaya and Keller (2026)*** showed that the functional influence of L5 on L2/3 rules out specific circuit motifs inspired by classical predictive coding, further supporting the idea that predictive representations and teaching signals are the main mode of communication between cortical areas rather than prediction errors.

However, once a suitable set of candidate circuit elements for encoder, integrator, and predictor are identified, the model leads to a concrete prediction. In particular, acute perturbations that disrupt the coupling between encoder and integrator components should primarily alter the representations formed by the integrator–predictor circuit, while leaving the activity patterns associated with the encoder largely preserved. By contrast, chronic disruptions of this coupling are expected to interfere with representation learning itself, preventing the gradual alignment of internal representations and the emergence of stable, abstract latent variables.

Beyond the neocortex, the proposed separation between encoding, integration, and prediction naturally invites comparison with hippocampal function. A substantial body of work has characterized the hippocampus as a predictive system that learns temporally extended representations of experience, often formalized in terms of successor representations or predictive maps of future states (***Stachenfeld et al., 2017***; ***Gershman, 2018***; ***Whittington et al., 2022***). In this view, hippocampal activity reflects expectations about future sensory or cognitive states rather than a reconstruction of current input, consistent with the representational emphasis of predictive learning. Recurrent dynamics within hippocampal circuits, particularly in CA3, together with sequential activity observed during navigation and replay, have been interpreted as supporting temporal integration and prediction (***Lisman, 1999***; ***Levenstein et al., 2024***). From this perspective, hippocampus and cortex may implement complementary predictive architectures operating at different temporal and spatial scales, with hippocampal representations providing rapidly learned, temporally extended predictions that could scaffold slower cortical representation learning (***Buckner, 2010***; ***Schapiro et al., 2017***). While we did not explicitly model hippocampal circuitry, RPL’s emphasis on internal prediction aligns naturally with theories that treat the hippocampus as a generator of predictive representations rather than a passive memory store (***Buzsáki and Tingley, 2018***).

Overall, our results suggest that experimental observations traditionally interpreted through the lens of hierarchical predictive coding may instead reflect a more distributed predictive architecture, in which different circuit elements specialize in complementary roles. JEPAs provide a useful conceptual scaffold for organizing these roles, while remaining compatible with multiple circuit layouts and known deviations from strictly hierarchical organization. The model presented in this article gives one possible instance, but it provides also a blueprint outlining which minimal computations must be present. Where these reside, and how precisely they are implemented remains an open question. Elucidating the architectural motifs the brain employs, and under what conditions, will require targeted experimental and theoretical work.

### Limitations and future research directions

Our work has several limitations that we plan to address in the future. First, the link between RPL and concrete circuits in the brain is under-constrained. One key simplification of our model is the lack of top-down connections. Concretely, we assumed that the predictor in the h-RPL model is entirely local, with no top-down input, a crude simplification of cortical anatomy where top-down connections are pervasive and considered essential to provide lower sensory areas with high-level abstract representations (***Vezoli et al., 2021***). Adding such connections would allow modulating predictions within each circuit with global context from higher-level circuits broadly consistent with experimental findings on the role of feedback in sensory cortices (***Han and Helmchen, 2024***; ***Richter et al., 2024***; ***Westerberg and Roelfsema, 2025***). A similar combination of a feedforward sensory pathway with a separate recurrent feedback pathway has been previously proposed as the “dual counterstream” architecture, an organizational principle of cortical structure (***Markov et al., 2013***). Exploring this connection and its functional implications is an exciting avenue for future research.

Another limitation of our model is that we relied on backprop, a biologically implausible training method. While we have shown that circuit-local training can be effective with h-RPL (***Figure 9***), we still used backprop, and more specifically backpropagation through time (BPTT), to train each circuit. In the future, we will investigate how biologically plausible credit assignment strategies could make the learning algorithm more credible (***Guerguiev et al., 2017***; ***Whittington and Bogacz, 2017***; ***Sacramento et al., 2018***; ***Lillicrap et al., 2020***; ***Payeur et al., 2021***; ***Laborieux and Zenke, 2022***; ***Rossbroich and Zenke, 2023***; ***Song et al., 2024***; ***Goemaere et al., 2025***). Another promising direction is to explore biologically plausible learning rules for recurrent architectures that approximate BPTT (***Murray, 2019***; ***Zenke and Ganguli, 2018***). This could involve analyzing RPL’s learning dynamics and factorizing BPTT into an eligibility trace and a learning signal, as in previous work (***Zenke and Ganguli, 2018***; ***Bellec et al., 2020***).

Moreover, latent-variable learning in RPL was sensitive to the prediction time horizon and the underlying distribution of the latent variable. In this article, we have only explored this connection to a limited extent (***Figure 5—figure Supplement 1***; ***Figure 5—figure Supplement 2***; Supplementary Table S4). A systematic investigation of how the time horizon and latent distribution influence the learned representations will be the goal of future research. Furthermore, future work will investigate the influence of stochasticity more systematically in the underlying latent dynamics during training and inference, while exploring improvements to RPL under non-deterministic conditions.

Finally, we relied on linear decodability as a proxy to quantify the quality of learned representations. To that end, we extended the commonly used linear classification strategy to linear regression of dynamic latent variables. While we saw that RPL embeds unrelated latent variables in orthogonal subspaces and improves overall linear decodability, a key question is why. In fact, there is no immediately apparent reason why the learned representations should be embedded on linear manifolds, and presumably they are not. One reason behind the improved linear decodability may be that linear decoding benefits from the orthogonal embedding of unrelated factors, combined with the high embedding dimension. An important question for future research is thus to gain a deeper theoretical understanding of how representations of continuous latent variables emerge in self-supervised learning JEPAs and which inductive biases lead to representations conducive for their linear decodability.

Extracting abstract representations of the world’s invariant and dynamic features is a crucial function of the brain that enables intelligent behavior. Here, we have presented a plausible computational model of self-supervised representation learning for both object identity and dynamic latent variables. Our framework extends a long tradition of predictive processing models while going beyond classical predictive coding and pixel space prediction. Specifically, our work builds a conceptual bridge between neuronal circuits and JEPAs, a promising machine learning architecture that does not rely on a reconstruction objective. While many important questions remain open, we believe that our model provides a fresh approach to understanding how the brain forms abstract internal representations from continuous sensory experience, and that exploring the research avenues outlined above holds great promise for gaining deeper insights into the computational principles underlying biological intelligence.

## Data and code availability

The code to reproduce all modeling results in this article is available on Github^***1***^. We further used the following publicly available datasets: The Librispeech dataset is freely available for download^***2***^. We used train and test splits and force-aligned phoneme annotations as defined previously (***Oord et al., 2018***) and made available by the authors^***3***^. The TLT12 dataset^***4***,***5***^ and generation code^***6***^ are freely available online. The mouse arena videos are available on Figshare^***7***^. Further, the two down-sampled videos used in this article are included in our Github repository. Finally, the plots for the macaque PFC experiment (***Bellet et al., 2024***) were reproduced using data^***8***^ and analysis code^***9***^ shared publicly by the authors.

## Acknowledgments

We thank Anna Vasilevskaya, Georg Keller, and all Zenke Lab members for inspiring discussions throughout this project. Special thanks to Peter Buttaroni for contributing illustrations for figures. This project was supported by the Swiss National Science Foundation (Grant Number PCEFP3_202981), EU’s Horizon Europe Research and Innovation Programme (Grant Agreement No. 101070374, CONVOLVE) funded through SERI (Ref. 1131–52302), and the Novartis Research Foundation.

## Author contributions

Ashena Gorgan Mohammadi & Manu Srinath Halvagal: Formal analysis, Investigation, Methodology, Software, Visualization, Writing – original draft, Writing – review and editing; Friedemann Zenke: Conceptualization, Formal analysis, Funding acquisition, Investigation, Methodology, Methodology, Software, Supervision, Writing – original draft, Writing – review and editing.

## Methods

In this article, we studied different representation learning paradigms that rely on prediction errors computed in internal representation space. We quantified their ability to extract latent variables like object identity, position, velocity, and orientation from streaming data such as video or audio. In all cases, we used deep neural networks comprised of three distinctly connected network elements: a feedforward encoder network, a recurrent integrator network, and a feedforward predictor network.

### Learning objectives

All networks were trained using backprop to optimize one of the following learning objectives defined on the internal representations.

#### Recurrent predictive learning (RPL)

While most predictive processing models optimize for reconstructing stimuli in input space, RPL optimizes a predictive objective function defined entirely on internal representations. Concretely, RPL is a particular JEPA which predicts the embeddings ***z****_t_*_+Δ_ of future stimuli using a predictor network *f* that acts on the representations ***c****_t_* from the integrator (cf. ***Figure 2a***). So, the RPL objective is defined as:

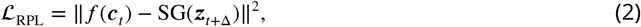

where SG corresponds to the Stop-Gradient operator. The RPL objective avoids representational collapse by combining the SG with an asymmetric prediction architecture and, consequently, does not require explicit variance regularization (***Chen and He, 2021***; ***Tian et al., 2021***; ***Srinath Halvagal et al., 2023***).

#### Invariance learning (IL)

Invariance learning is an alternative approach for representation-space predictive learning that uses a symmetric prediction scheme with explicit regularization terms (***Bardes et al., 2022***; ***Halvagal and Zenke, 2023***) for preventing collapse. This is also sometimes referred to as a joint-embedding architecture (***LeCun, 2022***; ***Assran et al., 2023***). The IL objective defined on the output of the predictor network in an architecture identical to the one used in RPL (cf. ***Eq. (2)***) is one such approach. We use this, along with a regularization objective *L*_reg_ similar to the one used by ***Halvagal and Zenke (2023)***, as a baseline for comparison with RPL. The overall IL objective function is then defined as:

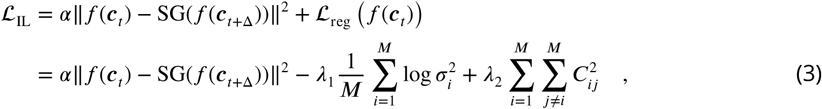

where *σ*_i_^2^ is the overall variance of the activity of neuron *i* in the output layer (i.e. *f*(***c****_t_*)), *C_ij_* is the correlation coefficient between the activities of neurons *i* and *j*, *α* is a constant that controls the strength of the invariance term and is set to 1 unless said otherwise, and *λ*_1_ and *λ*_2_ are constants that control the strength of each regularization term. The two terms in *L*_reg_ prevent the neurons from activity collapse and learning identical information, respectively. Note that the statistics in *L*_reg_ are calculated along both the time and batch dimensions.

Other variations of the IL objective can be defined to enforce invariance on the embeddings ***z****_t_* or representations ***c****_t_* of the RPL network. Mathematically, the objective functions corresponding to these variations can be written as:

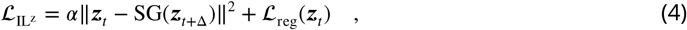

and

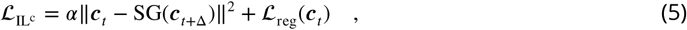

respectively with the target for *L*_reg_ changed in each case as indicated.

#### Context-predictive learning (CL)

Finally, the RPL objective is not the only possible implementation of a JEPA given our three-component network architecture. Alternatively, one could also optimize the same network for predicting future representations ***c****_t_*_+Δ_ from the integrator instead of ***z****_t_*_+Δ_, yielding the context-predictive learning (CL) objective function:

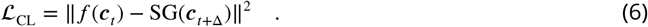

#### Additional variance-invariance-covariance (VIC) regularization

In networks with VIC regularization, we added the following regularization terms

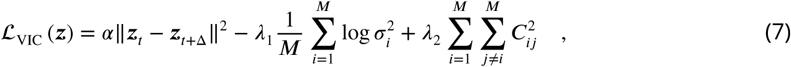

on the encoder’s embeddings, where *σ* and *C* are variance and covariance matrix of ***z***, to promote learning of rotation- and light-invariant features in the TLT12 dataset. More explicitly, RPL^VIC^ and IL with VIC regularization (IL^VIC^) objective functions are defined by

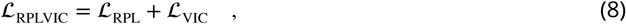

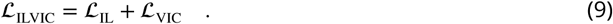

Note that *L*_VIC_ is similar to the IL*^z^* loss, except that the invariance term is fully symmetric, i.e., it does not include SG.

### Datasets and network models

We investigated RPL’s performance on multiple synthetic and real-world video and audio datasets with corresponding deep neural network architectures for the encoder, integrator, and predictor.

#### Synthetic moving animal video dataset

We generated the synthetic 32-frame moving animal videos by first generating sequences of the continuous latent variables. To that end, we randomly sampled an animal category and simulated its motion with a random initial velocity *v*_1_, *v*_2_ ∼ *Ѵ* (−8, 8) and rotation with a constant randomly sampled angular velocity 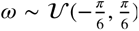. We also simulated motion in a third depth dimension by changing the scale of the animal starting with a random initial velocity *v*_3_ ∼ *Ѵ* (−0.125, 0.125). Furthermore, we assumed elastic collisions at the position boundaries in all three dimensions, which changed the direction of motion. Overall, the latent variables included the animal category *S*, position (*r*_1_, *r*_2_), scale *r*_3_, velocity ***v*** = (*v*_1_, *v*_2_, *v*_3_), planar orientation *θ*, and the corresponding angular velocity *ω*. We sampled the initial orientation from a uniform distribution *θ* ∼ *Ѵ* (0, 2*π*). The initial position in the *r*_1_-*r*_2_ plane was sampled from a uniform distribution: *r*_1_, *r*_2_ ∼ *Ѵ* (8, 56), where 8 is the padding around the image, ensuring that the animal remained visible within the frame. The scale *r*_3_ was randomly initialized from and remained limited to the interval [0.2, 1].

We then sampled the latent variable trajectory at discrete intervals and constructed the corresponding 64 × 64 RGB video frames. Each frame contained one antialiased animal image scaled and rendered on a gray background. Here, a scale *r*_3_ = 1 corresponded to an image size of 32 × 32 pixels. Finally, we added independent Gaussian noise ∼ *N* (0, 0.1) to each pixel and color channel (***Figure 2—figure Supplement 1***). Using this procedure, we built a fixed corpus of 4000 32-frame videos for validation and testing, and randomly generated training sets of 16000 videos for each network model we trained.

#### Network architecture and training

We used a convolutional neural network (CNN) with six convolutional layers for the encoder, each with 32 filters of size 5 × 5 and padding of size 2, followed by a batch normalization layer and a rectified linear unit (ReLU) activation function. We used a stride of two for the first two layers to reduce the spatial dimensions, and stride one for later layers. The output of the last convolutional layer was flattened to obtain the embedding vector ***z****_t_*. This embedding was then fed to the integrator with a single layer of 512 long short-term memory (LSTM) units to obtain the internal representation ***c****_t_*. Finally, this representation was passed through the predictor, a feedforward Multi-Layer Perceptron (MLP) network with one hidden layer of 512 units and the ReLU activation function, followed by a linear output layer. We trained the networks using the AdamW optimizer (***Loshchilov and Hutter, 2019***) with learning rate 3 × 10^−4^, weight decay 10^−5^, and batch size 128 for 1000 epochs using the aforementioned objective functions with Δ = 1 on the 16000 training videos, i.e., a total of 512000 frames, using a cosine learning rate schedule. In the case of RPL and CL (***Eqs. (2)*** and (6)), we used a learning rate 10 times higher for the predictor network relative to the encoder, an established protocol for JEPA training (***Chen and He, 2021***). For IL variants, we used *λ*_1_ = 20 and *λ*_2_ = 200 for the regularization strengths.

To compare our model to established SSL, albeit not necessarily biologically plausible algorithms, we trained the same network architecture via an InfoNCE loss (***Oord et al., 2018***) with 10 negative samples per positive sample and using either a nonlinear (i.e., an identical MLP predictor to RPL) or linear predictor network (Supplementary Table S1).

#### Evaluation

We then evaluated the trained network models using linear probing as follows. First, we froze all network parameters and computed the internal representations ***c*** of the training and testing datasets. We then standardized these representations to zero mean and unit variance. We used the resulting representations of the training dataset to fit an ordinary least square linear regression model for each continuous latent variable and a linear classifier for animal category. Since orientation is not a linear variable, we used *a* = sin *θ* and *b* = cos *θ* as target variables for the linear readout, which was trained by minimizing the mean-squared-error. We additionally optimized a phase offset for each animal category to correct for orientation offset biases per animal in the shared representation space. Moreover, we decoded the angular speed |*ω*| from time-averaged video representations. We trained the linear classifier on time-averaged video frames of the training set using the AdamW optimizer with a learning rate of 10^−3^, weight decay of 10^−5^, and a batch size of 128 for 250 epochs. Throughout the article, we reported the classification accuracies and coefficient of determination *R*^2^ for regression on the held-out test data. Finally, to provide a lower reference value for comparison, we evaluated the readout performance as described above for an untrained, randomly initialized network as a control. Similarly, to compare with supervised learning, we also trained the same network end-to-end using the regression and classification readouts as supervised training objectives for optimizing the encoder and integrator. Note that the predictor is unused during evaluation in all cases, and also during supervised training.

#### Internal simulation of trajectories

To investigate the model’s ability to accurately simulate trajectories soley based on internal representations, we primed the network with the first eight frames of a video. After the initial priming phase, the predicted embeddings were fed back to the integrator autoregressively for the following sixteen time steps thereby generating simulated internal representations (***Figure 3a***). To evaluate the quality of the simulation, we then estimated the latent variables from the internal representations using the linear decoders described above and compared them to the actual latent variables of the ground truth trajectory (cf. ***Figure 3*** and ***Figure 3—figure Supplement 1***).

### Temporal Latent Traversal Twelve (TLT12)

To test RPL on stimuli with more realistic 3D motion, changing lighting conditions, and a larger number of latent variables, we generated the TLT12 dataset. The dataset consists of 120,000 32-frame videos featuring 120 3D objects across 12 categories, or synsets, from the ShapeNetCore database (***Chang et al., 2015***), 10 objects per category, with 1,000 videos per object, rendered in a box with varying floor hue and spot lighting conditions. We picked objects from the following categories: airplane, cap, car, chair, earphone, guitar, laptop, motorbike, mug, piano, telephone, and watercraft. Each image is characterized by the object category, its position ***r*** = (*r*_1_, *r*_2_, *r*_3_) in Cartesian coordinates and orientation ***θ*** = (*θ*_1_, *θ*_2_, *θ*_3_) in Euler angles, the spot’s position (*θ*^spot^, *ϕ*^spot^) in spherical coordinates and its hue, and floor hue (***Figure 4a***). The object position is bounded to [−0.5, 0.5] along all axes, while the orientation is bounded to values within 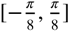 only along the first two axes. Additionally, *θ*^spot^ is restricted to values with 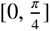.

We first generated smooth trajectories over the latent variables. To that end, we uniformly sampled initial values for each of the latent variables described above. We then randomly decided for each factor, with a probability of 0.8, whether they were allowed to change within a given sequence. The so determined *dynamic* factors were assigned a nonzero rate of change, initially sampled from a uniform distribution. We denote the rates of change, i.e., velocities for object translation and rotation, as ***v*** = (*v*_1_, *v*_2_, *v*_3_) and ***ω*** = (*ω*_1_, *ω*_2_, *ω*_3_) respectively, and use Δ for other variables. Further, we allowed the corresponding velocities of the dynamic factors to change following an Ornstein-Uhlenbeck process with a momentum of 0.95 sampled at discrete time intervals. For latent variables with bounded values, such as position, we used elastic collisions reflecting the latent variable within the boundaries by reversing its direction upon crossing the boundaries.

Following the above procedure we generated 1000 sequences for each of the 120 objects which were then rendered to 128 × 128 RGB images using custom written code based on work by ***Garrido et al. (2023)*** and BlenderProc (***Denninger et al., 2023***), leading to a total of 3,480,000 images. This step took approximately 500 GPU hours on cluster consisting of Nvidia V100 and A100 nodes. We then shuffled the samples and split the data at the sequence level in training, validation, and test sets with an 80:10:10 ratio. Thus different splits contain rendered images of the same objects. To also assess out-of-distribution generalization, we generated a separate test dataset following the above procedure but using ten different held-out objects for each category. The final dataset splits packaged together with the corresponding ground-truth factors using WebDataset format for ease of loading.

#### Network architecture and training

We used a variant of the ResNet-18 architecture (***He et al., 2016***) as the encoder, with half the number of filters in each layer and 3 × 3 filters in the stem. We removed the final fully connected layer, resulting in 256-dimensional embeddings after average pooling. The resulting embeddings were then fed to an integrator with a single layer of 256 LSTM units to generate the internal representations. Finally, the internal representations were fed to an MLP predictor with a single hidden layer of 1024 units and the ReLU activation function, followed by a linear output layer of 256 units to match the embeddings’ dimension. We trained *n* = 5 networks using the AdamW optimizer with learning rate 3 × 10^−4^, weight decay 10^−5^, batch size 128, and a cosine learning rate scheduler using RPL and IL objectives above with and without the additional VIC regularization, and with Δ = 2. We trained RPL^VIC^ and IL^VIC^ for 20 epochs, and RPL and IL for 50 epochs. For RPL and RPL^VIC^, we used a learning rate 10 times higher for the predictor network relative to the encoder. RPL^VIC^’s hyperparameters were selected by a Bayesian hyperparameter sweep to *α* = 0.03, *λ*_1_ = 0.9, and *λ*_2_ = 18. For IL and IL^VIC^, we used *α* = 13.08, *λ*_1_ = 39.16, and *λ*_2_ = 83.46, which were also determined through a Bayesian hyper-parameter sweep. We used the same hyper-parameters for the VIC regularization in IL^VIC^ for comparability.

#### Evaluation

We evaluated the quality of the learned representations using linear probing. To that end, we first extracted the internal representations ***c*** of all trained networks on the different subsets of the dataset. We then fitted logistic regression (using the LBFGS algorithm with a maximum of 1000 iterations) and ridge regression models with a regularization coefficient of 1.0 on 10% of the training set, and evaluated on the test set. Similar to the moving animal video dataset, we used sin and cos of variables with circular values as the regression target. To account for phase-offset differences across categories when estimating *θ*_3_ from the learned representations, we formulated phase-corrected ridge regression as a bilinear optimization problem, enabling a faster evaluation (see Supplementary Note S3 for details). We additionally trained *n* = 5 encoder-integrator networks end-to-end with a multi-task supervised training using AdamW optimizer with learning rate 10^−3^, weight decay 10^−5^, and batch size 128 for 50 epochs, following the same evaluation procedure on the internal representations. We also followed the same evaluation procedure on representations from *n* = 5 randomly initialized networks to provide a lower bound for downstream performance. To assess the power of the applied VIC on the encoder alone, we also trained *n* = 4 encoder networks with the same hyperparameters as RPL^VIC^ and followed the same evaluation procedure but on the encoder embeddings (Supplementary Table S2, VIC (enc)). However, for better comparability, we also fed the embeddings to randomly initialized integrators and evaluated the resulting internal representations (Supplementary Table S2, VIC). We additionally trained separate per-category ridge regression models with the same regularization coefficient on the representations of RPL^VIC^ and IL networks (cf. ***Figure 4e***). After removing all representations corresponding to the object class “earphones,” we trained another set of ridge regression models with the same regularization coefficient on orientation and rotation velocity factors, using the same phase-corrected rotation probe for *θ*_3_ as described above. The same trained regression models were later used to evaluate the representations extracted from the out-of-distribution test set (cf. ***Figure 4—figure Supplement 6***).

### Real-world videos of behaving mice

To study RPL’s ability to extract latent variables from real-world data, we used videos from the open source dataset of behaving mice in an open-field arena provided by ***Luxem et al. (2022)***. Concretely, we used two one-hour videos with a 25 Hz frame rate resulting in a total of 2 ×90000 frames. To split the data into training and test sets, we sliced the videos into one-minute segments from which we used the first fifty seconds for training and the last 10 seconds for testing. We downscaled all frames to 96 × 96 pixel frames using HandBrake (***Petit, 2003***) and subsampled the video segments at 5 Hz, i.e., only using every fifth frame, to decrease computational training cost. We further computed the proxy latent variables velocity, orientation, and position based on the dataset’s DeepLabCut (***Mathis et al., 2018***) estimates of tail root and nose position. To reduce label noise in the velocity estimates of each sample (***Figure 5—figure Supplement 1***), we smoothed them using a cumulative summation over time before standardizing the values. We further noted that the dataset contained erroneous nose position outside of the arena, for instance, whenever the nose was not visible when the animal was rearing. To prevent this label noise from entering our orientation estimates, we replaced the affected data points with linearly interpolated values.

#### Network architecture and training

We used a CNN with six convolutional layers, each with 32 filters as the encoder. We used 5 × 5 filters with padding and strides of size two for the first three layers, and 3 × 3 filters with padding and strides of size one for the following three layers. All convolutional layers were followed by a batch normalization layer and ReLU activation functions. The output of the last layer was flattened to obtain the embedding vector ***z****_t_*. This embedding was then fed to the integrator consisting of a single layer of 256 LSTM units to obtain the internal representation ***c****_t_*. Finally, the internal representations passed through an MLP predictor network with a single hidden layer comprised of 256 units followed by ReLU activation functions and a dense linear output layer. We trained the networks using the AdamW optimizer with learning rate 3 × 10^−4^, weight decay 10^−5^, and batch size 128 for 500 epochs using the aforementioned objective functions with Δ = 1 on the training set. In the case of RPL (***Eq. (2)***), we trained the predictor network with a 10 times faster learning rate than the rest of the network. When training using an IL objective, we additionally set the regularization strengths to *λ*_1_ = 20 and *λ*_2_ = 200.

#### Evaluation

We evaluated the networks on the test subset of the dataset using linear least squares with L2 regularization, i.e., ridge regression, with regularization coefficient of 1.0 on the trained internal representations ***c*** to decode the latent variables ***r****_N_*, ***v****_T_*, and *θ*.

### Real-world natural speech recordings

To test our representation learning framework in a different data domain, we used a 100-hour audio recording subset of the Librispeech dataset (***Panayotov et al., 2015***) for training and evaluating the network. Librispeech contains English audio book files from 251 speakers, with 41 force-aligned phoneme labels. The audio files were sampled at 16 kHz and there is a phoneme label for roughly every 10 ms of audio.

#### Network architecture and training

Following the work of ***Oord et al. (2018)***, we used a 1D CNN architecture acting on the raw audio waveform with 512 output units as the encoder. The resulting embeddings were then fed into the integrator, a single LSTM layer with 256 units, to generate the internal representations. The representations were then passed through an MLP predictor network with a single hidden layer of 512 units and ReLU activation function, followed by a dense linear output layer. The network was trained using the AdamW optimizer with learning rate 3 × 10^−4^, no weight decay, and batch size 256 for 500 epochs using the different objective functions with Δ = 8. The hyperparameter Δ was determined via grid search to be the optimal prediction horizon for RPL. In the case of IL objectives, we used *λ*_1_ = 2 and *λ*_2_ = 20 as regularization strengths.

#### Evaluation

We evaluated the trained networks on the test set following the same steps as above and by training linear readouts to decode the phoneme and speaker labels. The linear readout contained 251 units for speaker classification and 41 units for phoneme classification. We trained the linear classifiers using the AdamW optimizer with learning rate 10^−3^, weight decay 10^−5^, and batch size 64 (for speaker identity) for 50 epochs. Following the same procedure as in ***Oord et al. (2018)***, we use a batch size of one for training readouts for the phoneme classification task, as the start and end point annotations for phoneme labels are not provided. As before, we evaluated a randomly initialized network for comparison.

### Synthetic sequences of handwritten digits

To test RPL on more abstract stimulus sequences, we generated synthetic sequences of handwritten digit images consisting of digit triplets separated by zeros (***Figure 6a***). We generated the digit sequences by sampling clusters, triplets and digits for 64 time steps according to a structured grammar (cf. ***Figure 6b***). Specifically, we clustered the digits into four groups: {1, 2, 3}, {4, 5, 6}, {7, 8, 9}, and {0}. The permutations of the digits within each of the first three groups constituted the triplets, making a total of 18 possible triplets. Each digit sequence consisted of such triplets interspersed with zeros, starting with a triplet drawn from a randomly chosen cluster. Each triplet was either followed by another random triplet from the same cluster with a probability of 0.8, or a zero with a probability of 0.2. After a zero, a cluster group was chosen uniformly at random to continue the sequence. Overall, the latent variables in this case were the digit identity, the cluster group, and the triplet identity. Given the generated sequence of digits, we finally sampled corresponding 28 × 28 grayscale images from the MNIST training and test sets for training and testing respectively. Using this approach, we generated 10000 training sequences and 10000 test sequences for each model.

#### Network architecture and training

For the encoder, we used an MLP with two hidden layers consisting of 32 units, each followed by batch normalization and ReLU activation functions. We fed the resulting embeddings into a single-layer recurrent neural network (RNN) integrator with 128 units to generate the internal representations. The representations were further passed through a feedforward MLP predictor network with a single hidden layer of 128 units and ReLU activation function, followed by a linear output layer. We trained all networks using the AdamW optimizer with a learning rate 3 × 10^−4^, weight decay 10^−3^, and batch size 64 for 200 epochs using the different objective functions with Δ = 1 on the training sequences. For IL, we used *λ*_1_ = 1 and *λ*_2_ = 10 for the regularization strength.

#### Evaluation

To assess the quality of the learned representations, we first froze the parameters of the trained networks. Then, we extracted the representations of the training and test sets and standardized them as before. We subsequently trained a set of linear classifiers on these representations to decode digit identity and cluster identity at each time step, and triplet identity after observing the first two digits of any triplet subsequence (cf. ***Figure 6b***). We trained all three linear classifiers for 50 epochs using AdamW with a learning rate of 10^−3^, weight decay of 10^−5^, and a batch size of 128. As before, we also computed performance estimates for a random untrained baseline network as well as end-to-end supervised training.

### Learning successor-like representations

To investigate whether RPL learns successor-like representations, we modeled a stimulus sequence A-B-C-D as in the experiment (***Ekman et al., 2023***) with one-hot vectors. Since the one-hot encoding of the inputs alleviated the need for learning stimulus representations, we replaced the encoder with an identity matrix. For simplicity, we used a vanilla RNN with four units as integrator, whose weights we initialized with an identity matrix, and a dense linear predictor whose weights we initialized with a Glorot uniform distribution (***Glorot and Bengio, 2010***). Following a similar experimental setup as ***Ekman et al. (2023)***, we trained *n* = 35 networks for 352 epochs on the A-B-C-D sequence using the RPL objective with a learning rate of 10^−3^ for the integrator and 0.01 for the predictor. Further, we used a weight decay of 10^−4^ for all the components.

We trained logistic regression models on the internal representations ***c*** of the full sequences to classify the sequence locations. We then sorted the integrator units based on their selectivity to sequence locations. We computed the average weight matrix of all logistic regression models and visualized it to see if it resembles a successor matrix (***Figure 7b***). To generate the partial sequences, we used zero input vectors for the non-stimulated sequence locations and the corresponding one-hot vector for the stimulated location. We then computed the activity at stimulated and non-stimulated locations for each partial sequence by recording the mean population activity of the integrator units at each sequence location and subtracting the population response to a zero input (***Figure 7d***). These values were averaged over all partial sequences for each stimulated and non-stimulated location to obtain the activity at individual predecessor, stimulated, and successor locations (***Figure 7c***).

### Learning abstract sequence representations from local-global oddballs

To study whether RPL learns abstract sequence representations similar to the primate PFC, we modeled the local-global oddball experiment conducted by ***Bellet et al. (2024)***. We first extracted images of the ten visual stimuli used in the experiment and downsampled them to 32 × 32 pixel color images. We then constructed the oddball sequences from ***Bellet et al. (2024)*** consisting of pairs of stimuli organized in blocks of twenty sequences (scaled down from the original block size of two hundred). Each sequence comprised four stimulus presentations, either repeating the same stimulus (xxxx or xx for short) or with a local deviant at the last position (xxxY or xY for short). Here, x stands for either of the two items in each pair of stimuli and y for the other, fixed within one block. The first five sequences in each block were consistently either xx or xY, setting the global standard or “context” for that block. The next fifteen sequences were randomly set to either global standards (80%) or rare global deviants (20%). The global standards were repeats of the standard sequence, i.e., xx in an xx block (xx xx) or xY in an xY block (xY xY). The global deviants differed from the standard sequence only in the last position (xY xx or xx xY). The full sequence of stimuli also contained blank frames, one between each pair of consecutive stimuli in a given sequence, and four between sequences in the same block. We generated 2000 such blocks with the stimulus pairs and contexts uniformly chosen at random for each block for training the network model, and 2000 more for testing.

#### Network architecture and training

Since the stimulus set consisted of only ten images, we used a simple MLP with two hidden layers and batch normalization for the encoder. The MLP took in the full vector of image pixels as input and had 64 units in each hidden layer and its output layer. We then fed the embedding from the output of the encoder to an integrator consisting of a single LSTM layer with 256 units to generate the internal representations. These were then passed through an MLP predictor with one hidden layer with 256 units. Both the encoder and predictor MLPs used the ReLU activation function. For training, each block of 20 sequence presentations was given as one training sample sequence. We trained the network using the AdamW optimizer with learning rate 3 × 10^−4^, batch size 128, and weight decay 10^−3^ for 2000 epochs using the RPL objective function with Δ = 1. As before, we also used a 10 times higher learning rate for the predictor.

#### Evaluation

For evaluating the trained network, we closely followed the analysis of PFC population recordings done by ***Bellet et al. (2024)***. This consisted of training linear decoders for serial sequence position, stimulus identity, global context and presence of local and global deviance on internal representations ***c*** as above, but with specific sequence subtypes used for training the readout (cf. ***Fig 3*** and Methods in ***Bellet et al. (2024)***) as summarized below:

##### Serial position

For sequence position, we trained binary classifiers using logistic regression for each position in the sequence using only xx trials for training. This yielded predicted probabilities of items 1 through 4 at each time point of a sequence, plotted for the held-out test set in ***Figure 8b***.

##### Stimulus identity

Stimulus identity was defined as a fixed binary label for the two items in each of the five stimulus pairs. We trained a binary classifier for decoding this binary label using only the first three stimuli in each sequence for training. The projections of the representations onto the weights found by the binary classifier are plotted in ***Figure 8c*** for the held-out test set of sequences.

##### Global context

For decoding the context, we first averaged the representations over the first three stimulus pairs in every sequence attaching a binary label corresponding to whether the global standard was a repeat sequence (xx) or a sequence with a local oddball (xY). As above, we trained a binary classifier with logistic regression and plotted the projections of the representations from held-out sequences onto the weights found by training the binary classifier.

##### Deviance

For decoding both global and local deviance, we used the representation corresponding to the last stimulus presentation to train a binary classifier with logistic regression. Then, we plotted the projections of the representations for the test set onto the weights of the binary classifier.

All linear decoders were trained with AdamW using the learning rate 10^−3^ and weight decay 10^−5^.

### Hierarchical RPL (h-RPL)

To assess the effectiveness of RPL in a hierarchical setting, we constructed a network from six RPL circuits each with a shallow single-layer 2D CNN with 32 filters of size 5 × 5 and padding of size 2, followed by a batch normalization layer and a ReLU activation function. The first two convolutional layers had a stride of two while the convolutional layers in higher circuits had a stride of 1. Note that this stack of shallow instantaneous encoders in h-RPL results in the same deep convolutional encoder used in RPL for the same dataset. The encoder output was sent to the next circuit in the hierarchy and a flattened version was sent to the local integrator circuit, an LSTM with 512 units (***Figure 9a***) for all circuits. Each circuit also had its own MLP predictor with a single hidden layer comprised of 512 units with ReLU activation functions and followed by a dense linear output. Importantly, the different circuits were gradient isolated and independently optimized the RPL objective (***Eq. (2)***) at each level of the hierarchy, without backpropagating the gradients between circuits. Finally, we trained the model on the synthetic moving animal videos (cf. ***Figure 2***) using the AdamW optimizer with learning rate 3 × 10^−4^, weight decay 10^−5^, and batch size 128 for 500 epochs using the RPL objective. As before, we evaluated the representations using linear probing albeit now for each circuit individually.

---

**Figure 2—figure supplement 1.**
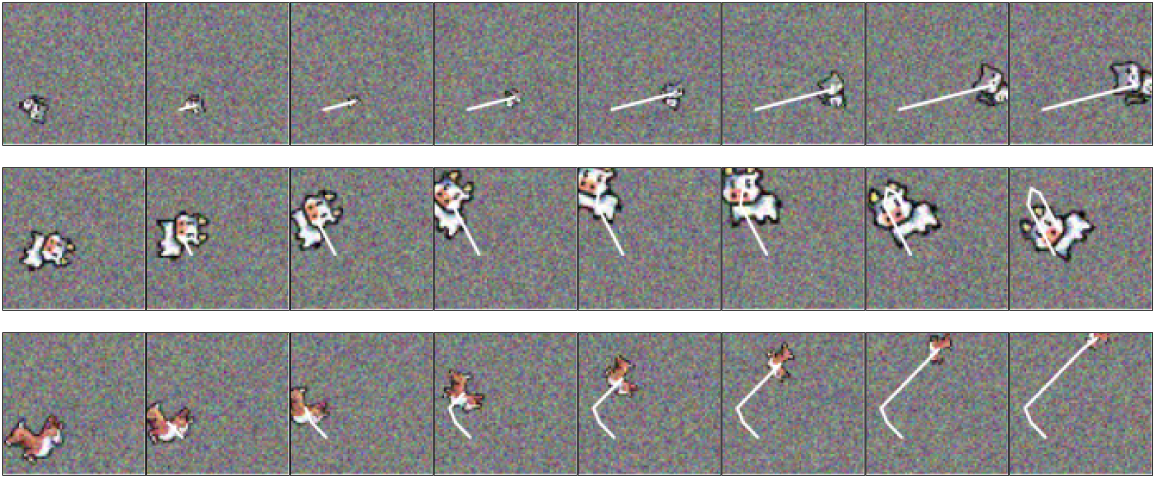
Sample video frames of the moving animals data input to the models. The white line shows the past motion trajectory of the animal and is invisible to the model.

**Figure 2—figure supplement 2.**
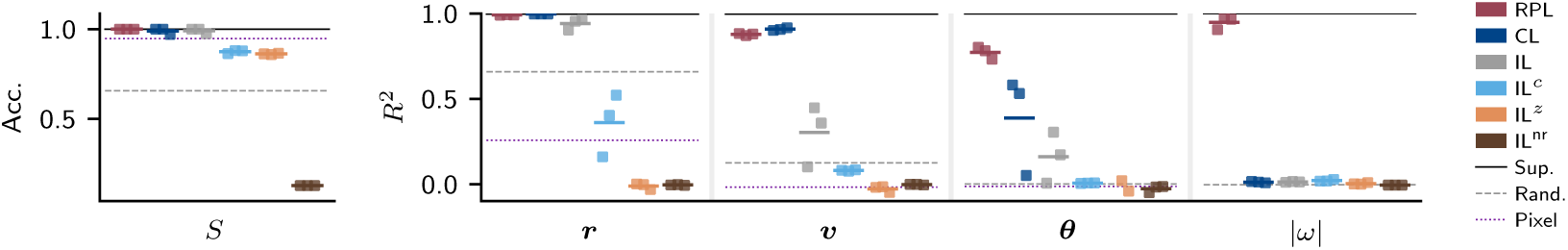
Comparison of representations learned from moving animal videos with different objectives. Linear readout accuracy of animal categories *S* and coefficient of determination (*R*^2^) (right) for position ***r***, velocity ***v***, angular speed (|*ω*|), and orientation (***θ***). RPL (red) (***Eq. (2)***) learns untangled representations of all latent variables. CL (dark blue) (***Eq. (6)***) fails to learn untangled representations of rotational dynamics. IL (gray) (***Eq. (3)***) learns untangled representations of animal category and position but it fails to learn other dynamic latent variables. IL objectives defined on the encoder (IL*^z^* (light blue); Eq.(4)) and the integrator (IL*^c^* (orange); Eq.(5)) are incapable of extracting untangled representations of all latent variables. Importantly, removing the regularization term from IL leads to fully collapsed representations (IL*^nr^* (brown)). Supervised training performance (solid black line), performance of training readouts on representations from randomly initialized network (dashed gray line), and performance on training directly on flattened pixel values (dotted purple) are provided as baselines.

**Figure 2—figure supplement 3.**
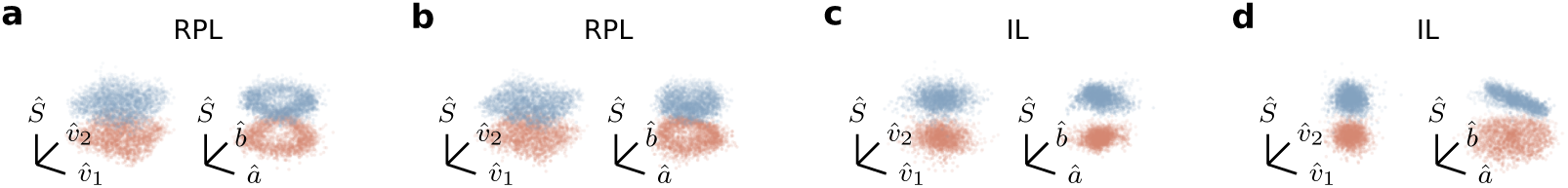
3D projections of learned representations of dog and cat planar velocity and orientation. **(a), (b)** Projections of learned representations by two other RPL models. **(c), (d)** Same for IL models.

**Figure 2—figure supplement 4.**
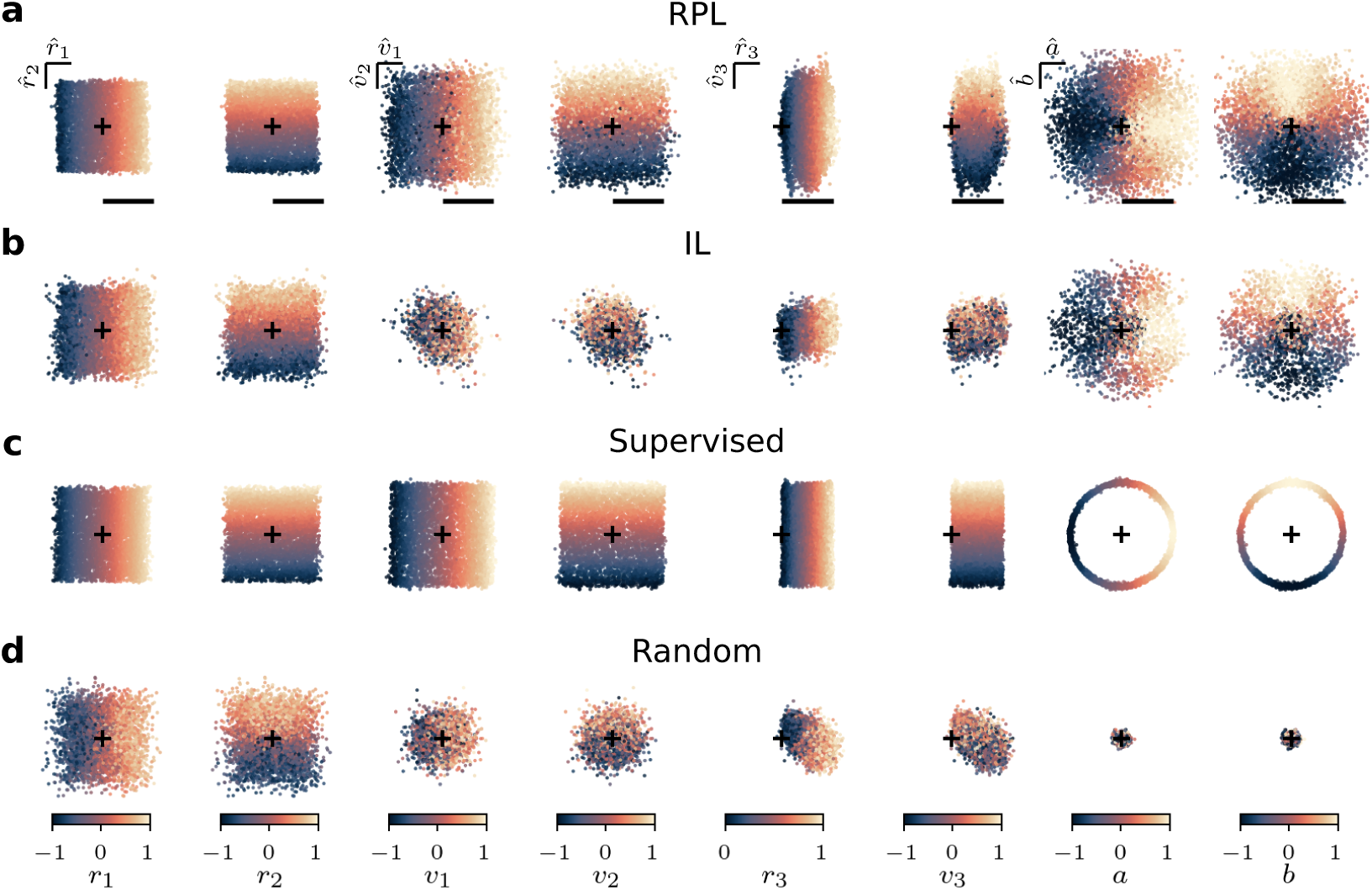
Projections of learned representations of motion factors from moving animal videos. **(a)** Projections of the representations learned with RPL along directions that are maximally predictive of pairs of latent factors (*r*_1_, *r*_2_), (*v*_1_, *v*_2_), (*r*_3_, *v*_3_) and (*a*, *b*). For each pair, the projections are plotted twice, colored by the true values of the first and second latent factor each time. **(b)** Same as (a) but for the representations learned with IL. **(c)** Same as (a) but for the representations learned with supervised training. **(d)** Same as (a) but for the network at random initialization.

**Figure 2—figure supplement 5.**
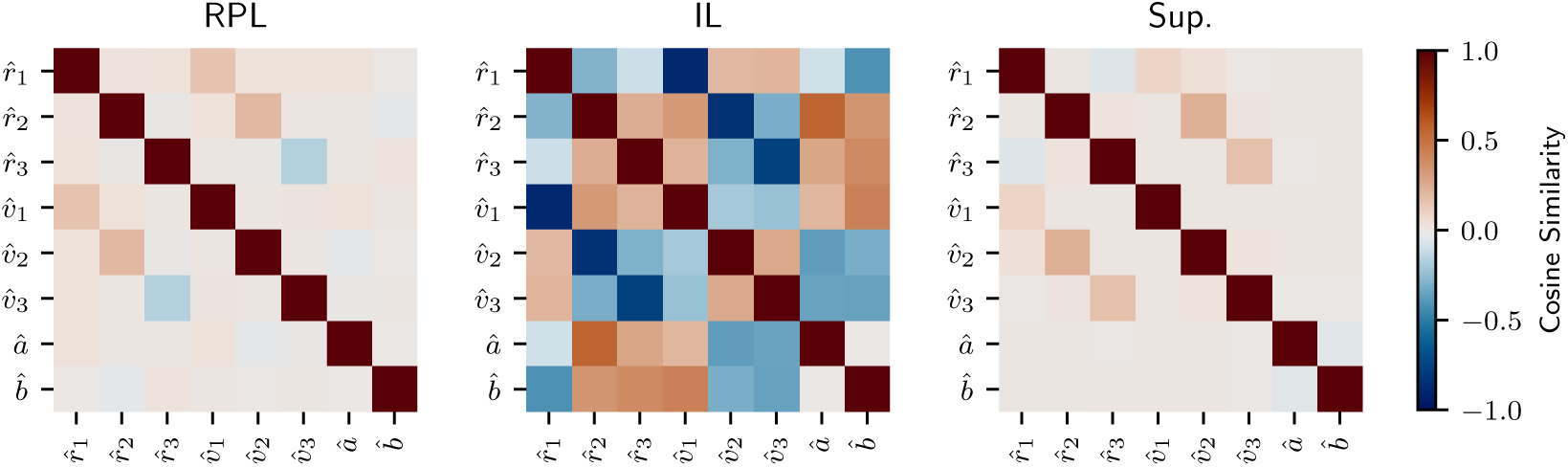
Orthogonality of learned representations of motion factors from moving animal videos. Pairwise cosine similarities between the maximally predictive directions of the learned representations for the dynamic latent variables.

**Figure 2—figure supplement 6.**
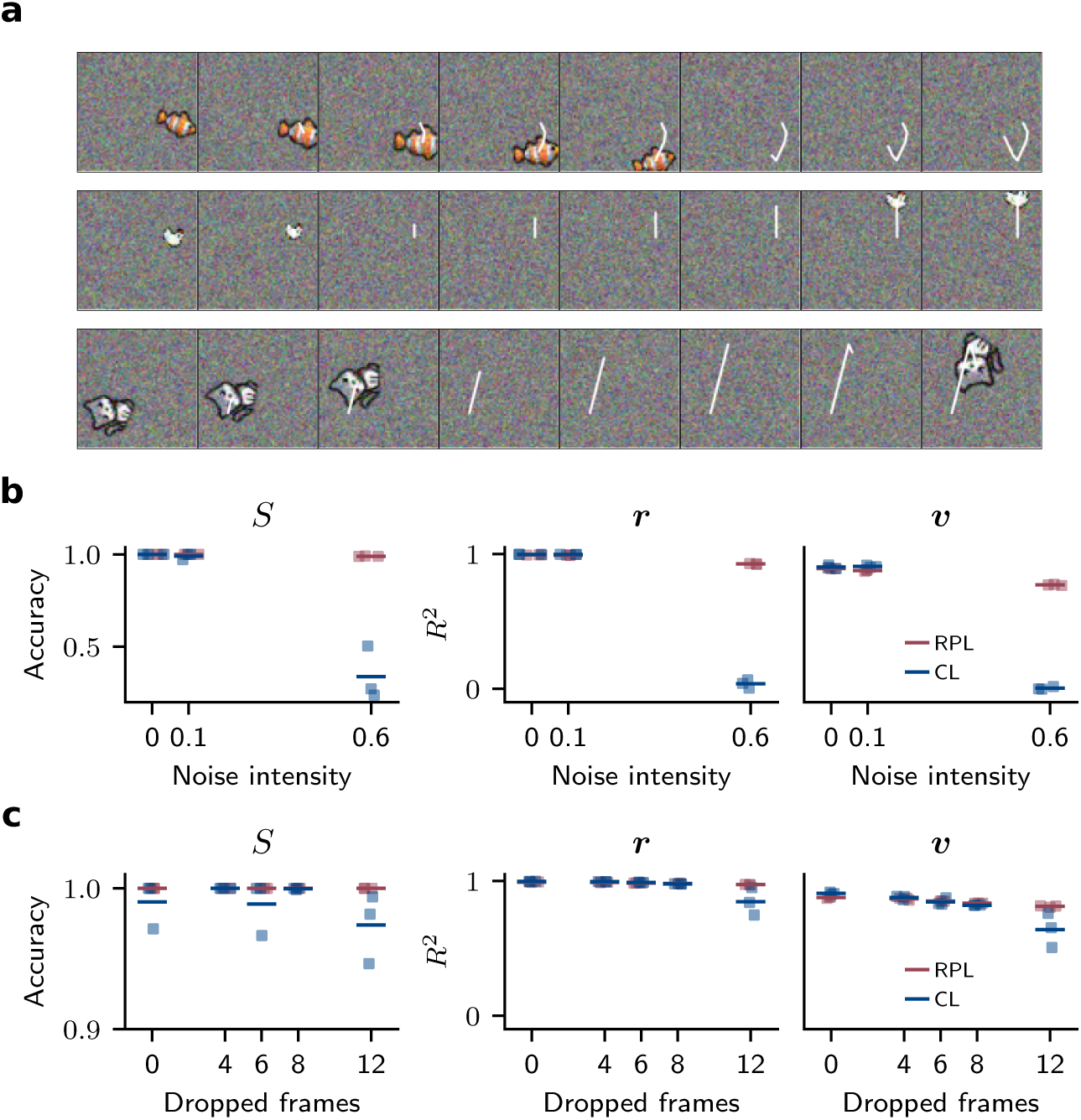
Effect of pixel noise and frame dropout on the learned representations from moving animal videos. **(a)** Sample video frames of the moving animals data with frame dropout. The white line shows the motion trajectory of the animal and is invisible to the model. **(a)** The accuracy of linear classifiers on animal categorization task (left) and coefficient of determination (*R*^2^) of linear regression on position (middle) and velocity (right) tasks for representations learned with RPL and CL are shown for different levels of Gaussian noise added independently to pixel values of each video frame. RPL is robust to high levels of noise, while CL is not. **(b)** Same as (a) but for frame dropout. While representations learned with both objectives have lower performance for larger numbers of dropped frames, the effect is more pronounced for CL than RPL.

**Figure 3—figure supplement 1.**
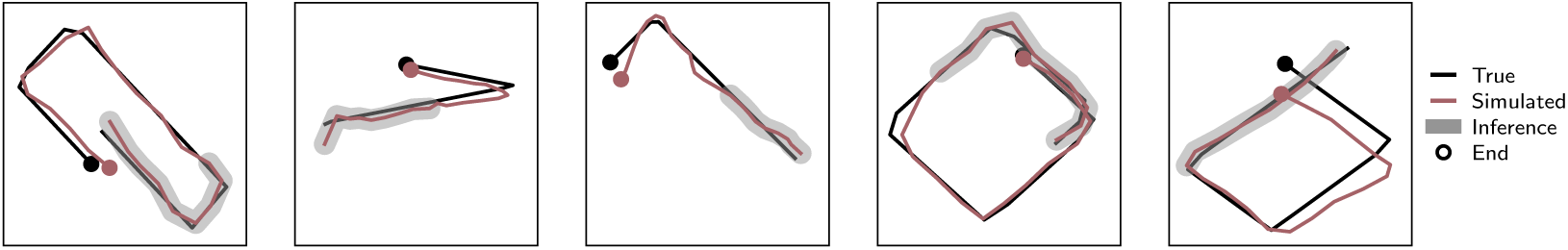
Simulated motion trajectories in the *r*_1∕2_ plane for moving animals data. More examples of simulated motion using the learned predictive representations.

**Figure 4—figure supplement 1.**
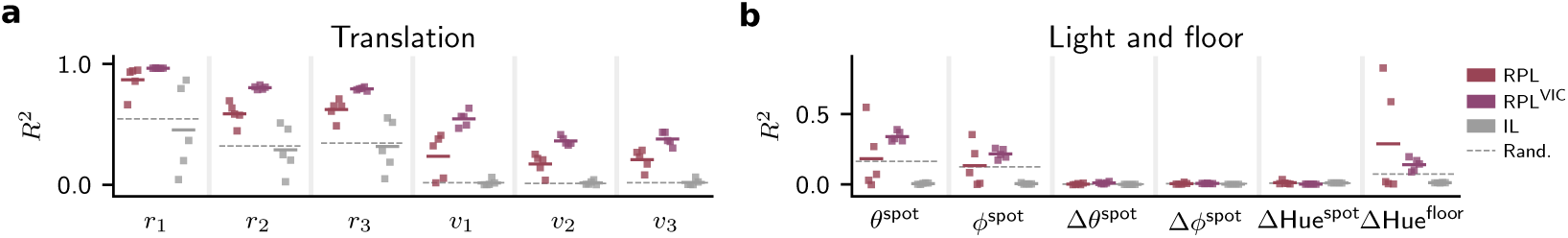
Linear decoding performance on TLT12 dataset for individual. (a) translation, and (b) light spot and floor factors.

**Figure 4—figure supplement 2.**
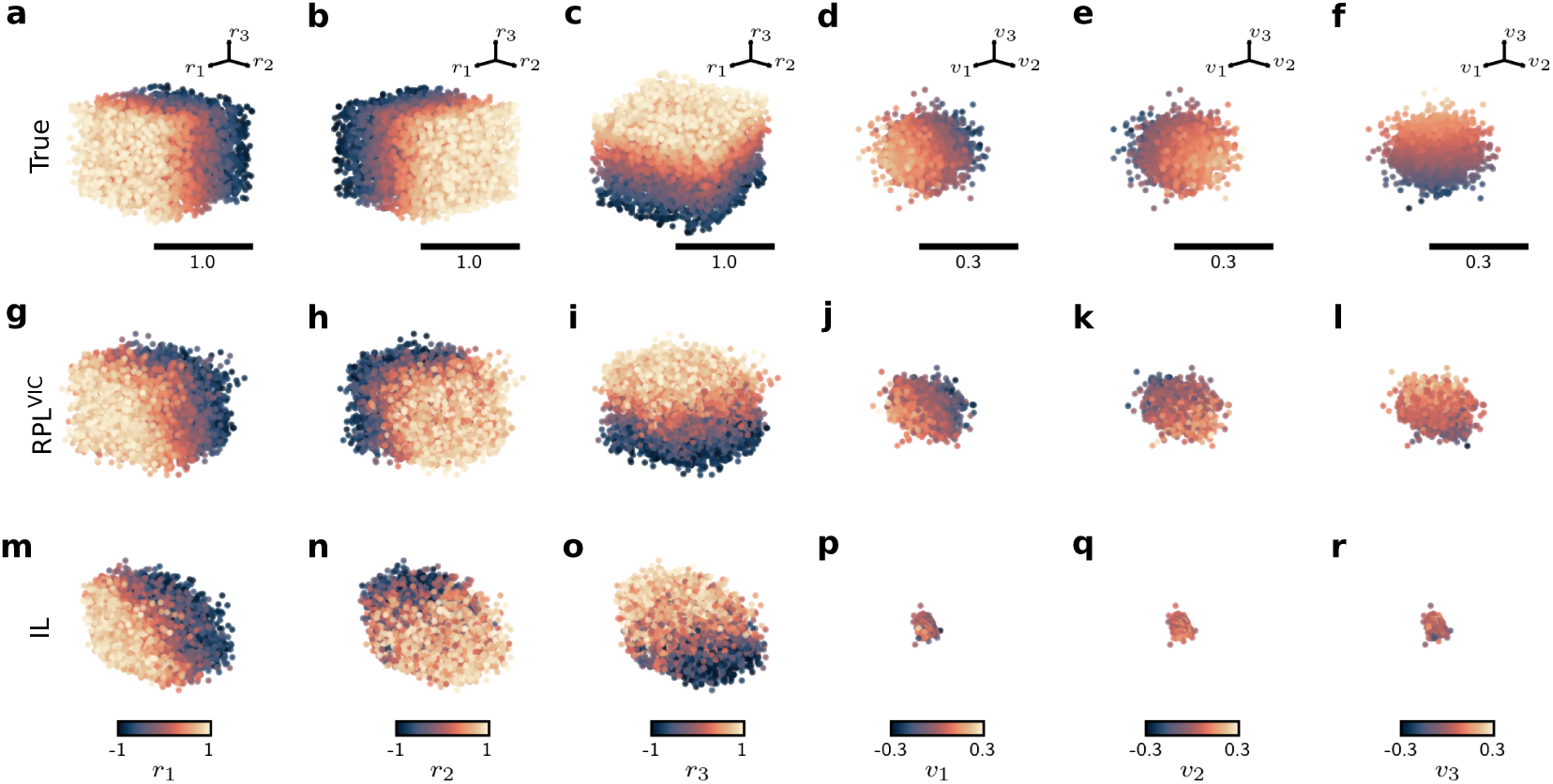
Linear projections of ground truth and learned representations of translation factors from TLT12. (**a–c)** Ground truth factor representations of object position in 3D space, colored by *r*_1_, *r*_2_, and *r*_3_ respectively. (**d–f**) Ground truth factor representations of object translation velocities colored by *v*_1_, *v*_2_, and *v*_3_ respectively. **(g–i)** Linear projections of representations from a trained RPL model onto directions coding for object position in 3D space, colored by *r*_1_, *r*_2_, and *r*_3_ respectively. **(j–l**) Linear projections of representations from a trained RPL model onto directions coding for object translation velocities in 3D space, colored by *v*_1_, *v*_2_, and *v*_3_ respectively. **(m–r)** Same as (**j–l)** but for representations from a trained IL model.

**Figure 4—figure supplement 3.**
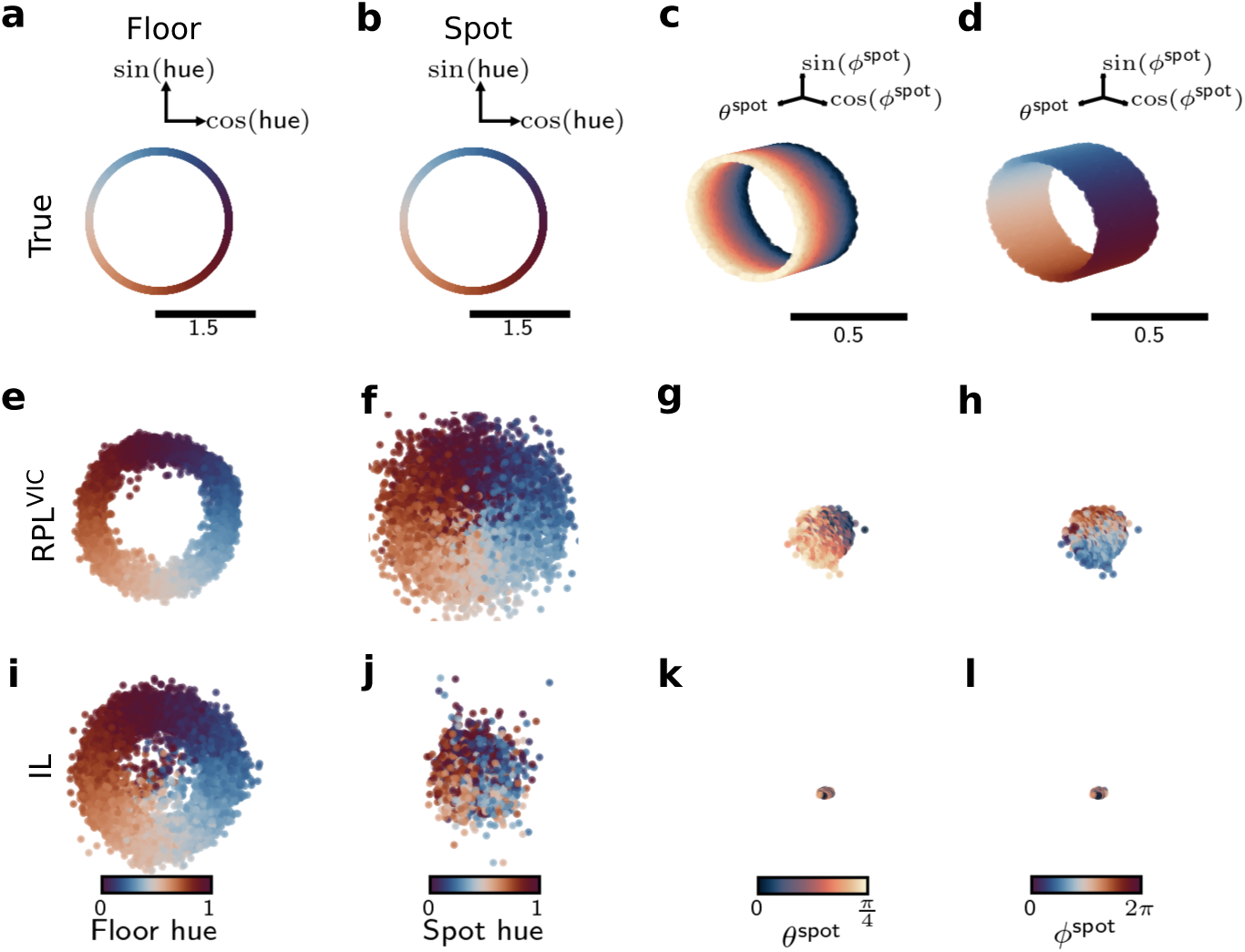
Linear projections of ground truth and learned representations of light spot and floor factors from TLT12. **(a), (b)** Ground truth factor representations of light spot elevation (*θ*^spot^) and azimuth (*ϕ*^spot^) colored by *θ*^spot^ and *ϕ*^spot^ respectively. (**c), (d)** Ground truth factor representations of light spot and floor hue respectively. (e) Linear projections of representations learned by RPL onto directions coding for *θ*^spot^ and *ϕ*^spot^ colored by *θ*^spot^. (f) Same as (e) but colored by *ϕ*^spot^. (g) Linear projections of representations learned by RPL onto directions coding for light spot hue. (h) Linear projections of representations learned by RPL onto directions coding for floor hue. (i)-(l) Same as (e)-(h) but for representations learned by IL.

**Figure 4—figure supplement 4.**
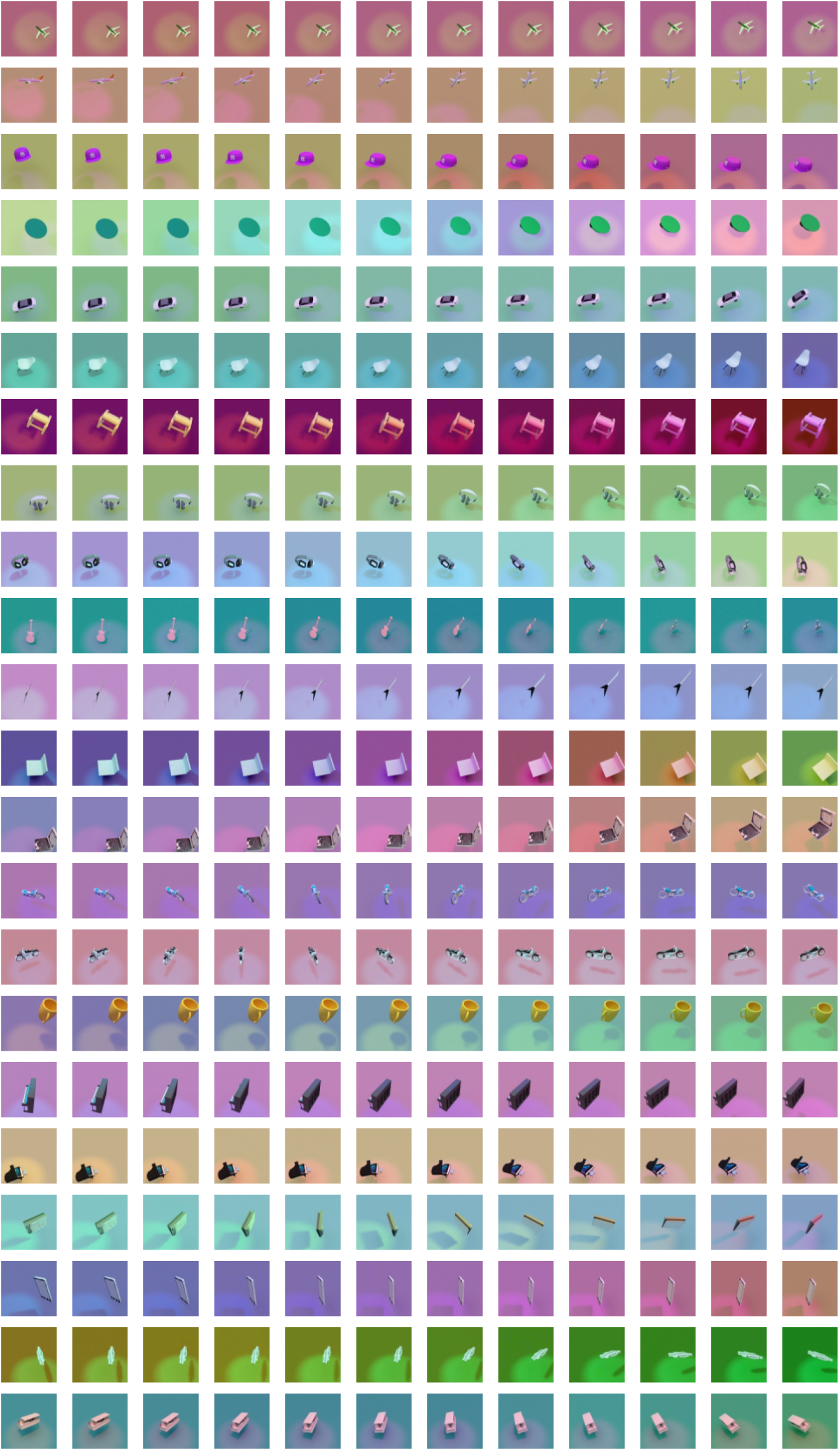
Sample video frames from TLT12. Each row demonstrates consecutive initial frames within a video sample.

**Figure 4—figure supplement 5.**
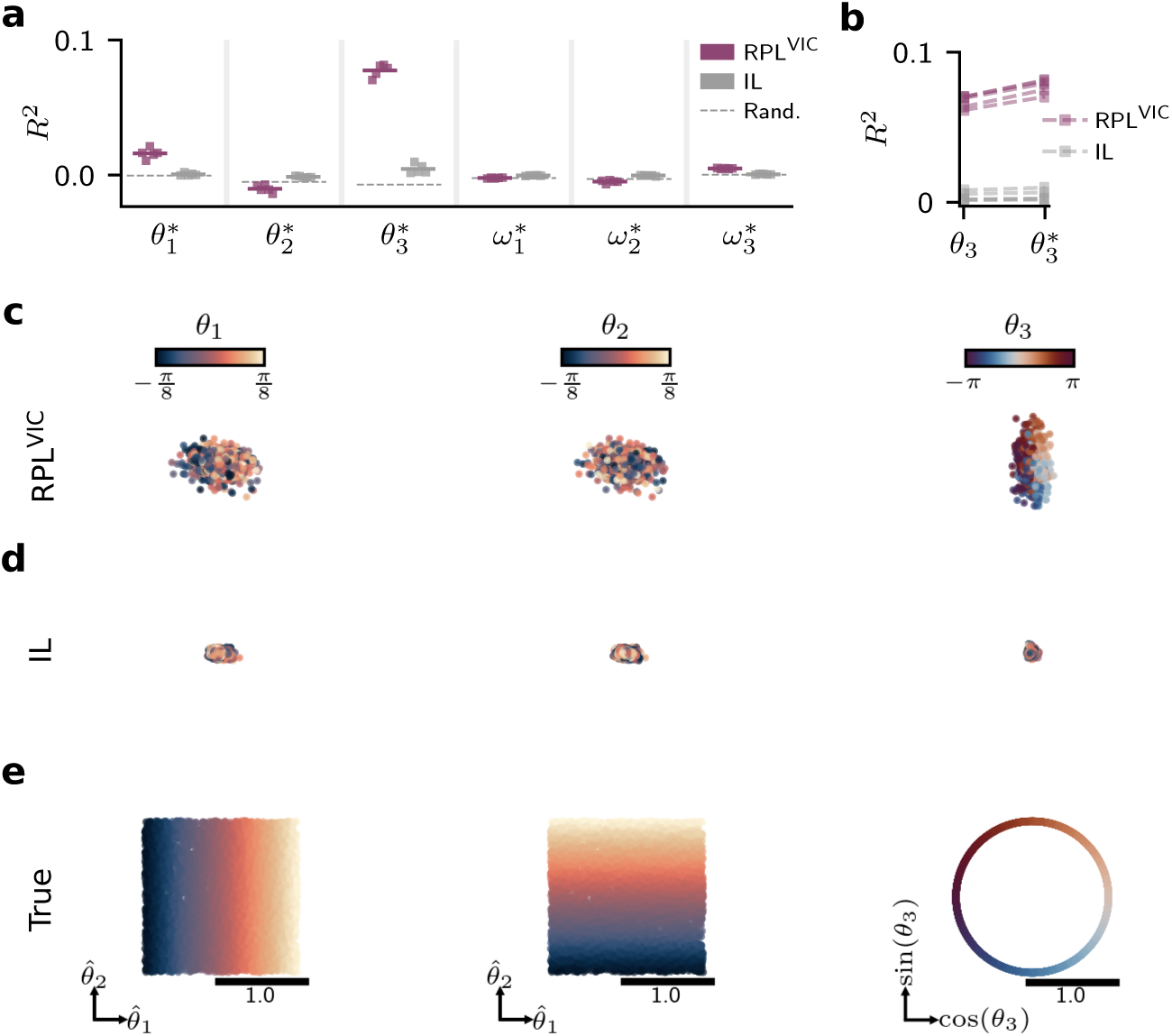
Orientation coding in representations learned by RPL and IL models from TLT12 dataset. **(a)** Coefficient of determination *R*^2^ of linear decoders on object orientation and rotation velocities in 3D space for representations learned with RPL (red) and IL (gray) models (*n* = 5). The decoders are trained on all objects but earphones, i.e. less symmetric objects along the third axis. The average performance of randomly initialized networks (gray dashed lines) is shown as a reference. **(b)** Decoding performance on the orientation along the third axis *θ*_3_ trained on all objects in comparison to decoding performance *θ*^∗^ after removing samples from earphone category for representations learned with RPL (red) and IL (gray) models. This result suggests the existence of a shared rotation coding axis along the third dimension. **(c)** Linear projections of representations from a trained RPL model onto directions coding for object rotation in 3D space, colored by *θ*_1_, *θ*_2_, and *θ*_3_ respectively. There is a clear distinction between negatively and positively oriented objects along the first axis and a relatively circular coding pattern along the third axis. **(d)** Same as (c) but for representations from a trained IL model. The results indicate representational collapse w.r.t. rotation factors. **(e)** Ground truth representations of the rotation factors, colored by *θ*_1_, *θ*_2_ and *θ*_3_ respectively.

**Figure 4—figure supplement 6.**
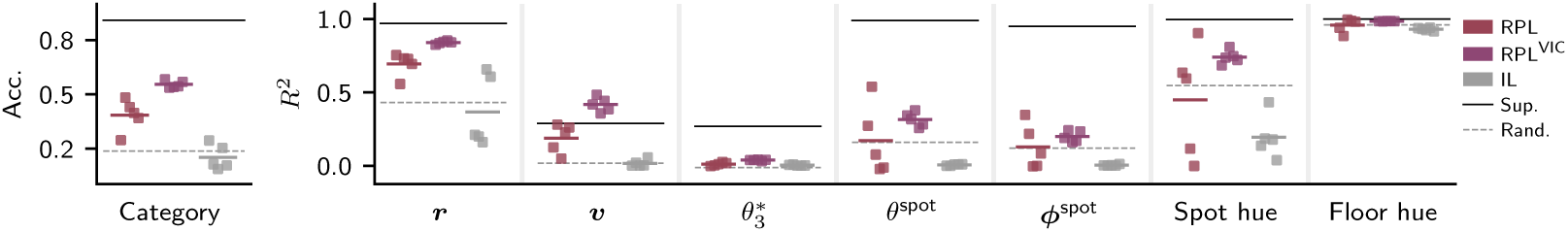
Linear decoding performance on representations of out-of-distribution test samples from TLT12 dataset extracted from different trained models. All models show a drop in classifying novel objects. This drop is less significant for RPL. However, all models represent the dynamic latent variables as reliably as they do for the unseen but known objects (cf. ***Figure 4c***), indicating that the representations are independent of individual objects.

**Figure 5—figure supplement 1.**
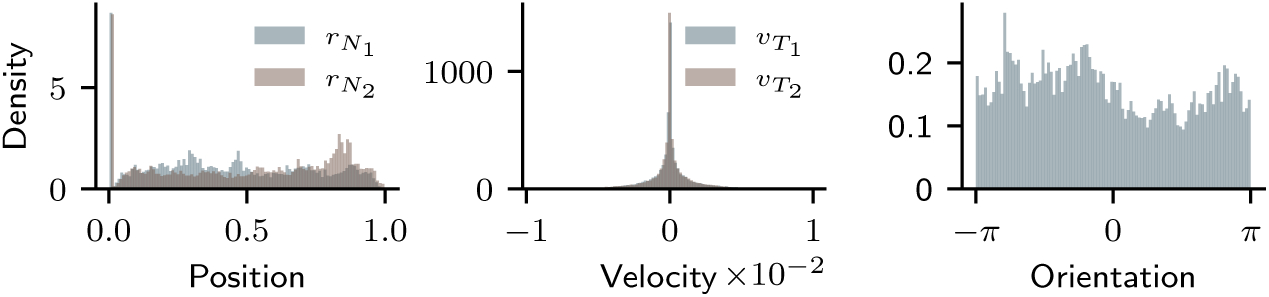
Distribution of latent variables in the training and test sets in the mouse data.

**Figure 5—figure supplement 2.**
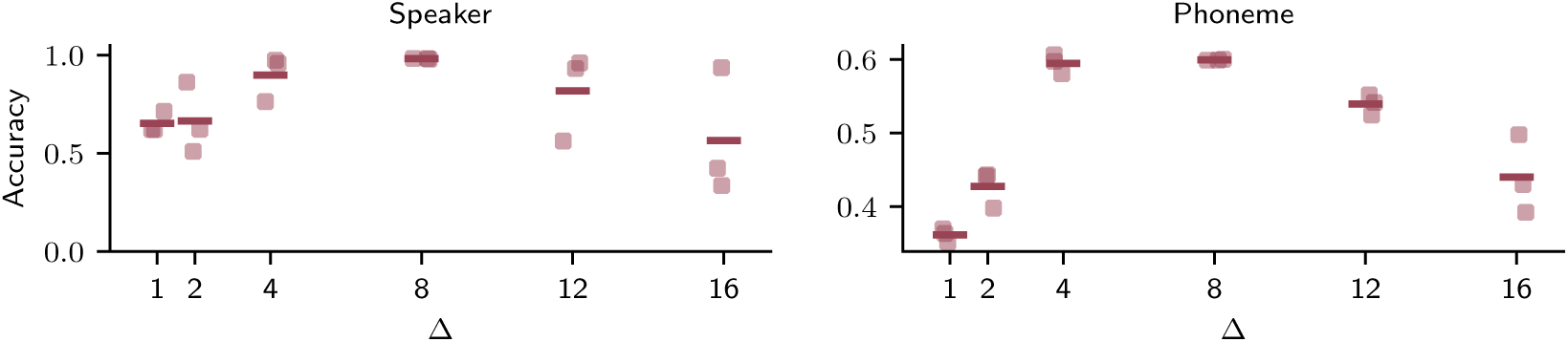
Downstream performance as a function of prediction horizon. Speaker and phoneme classification accuracy as a function of prediction horizon (Δ) in RPL in natural speech recordings. The optimal performance in both tasks is achieved with a Δ = 8 which roughly corresponds to average phoneme length (∼ 80 ms). This result highlights the role of prediction horizon in learning useful representations.

**Figure 6—figure supplement 1.**
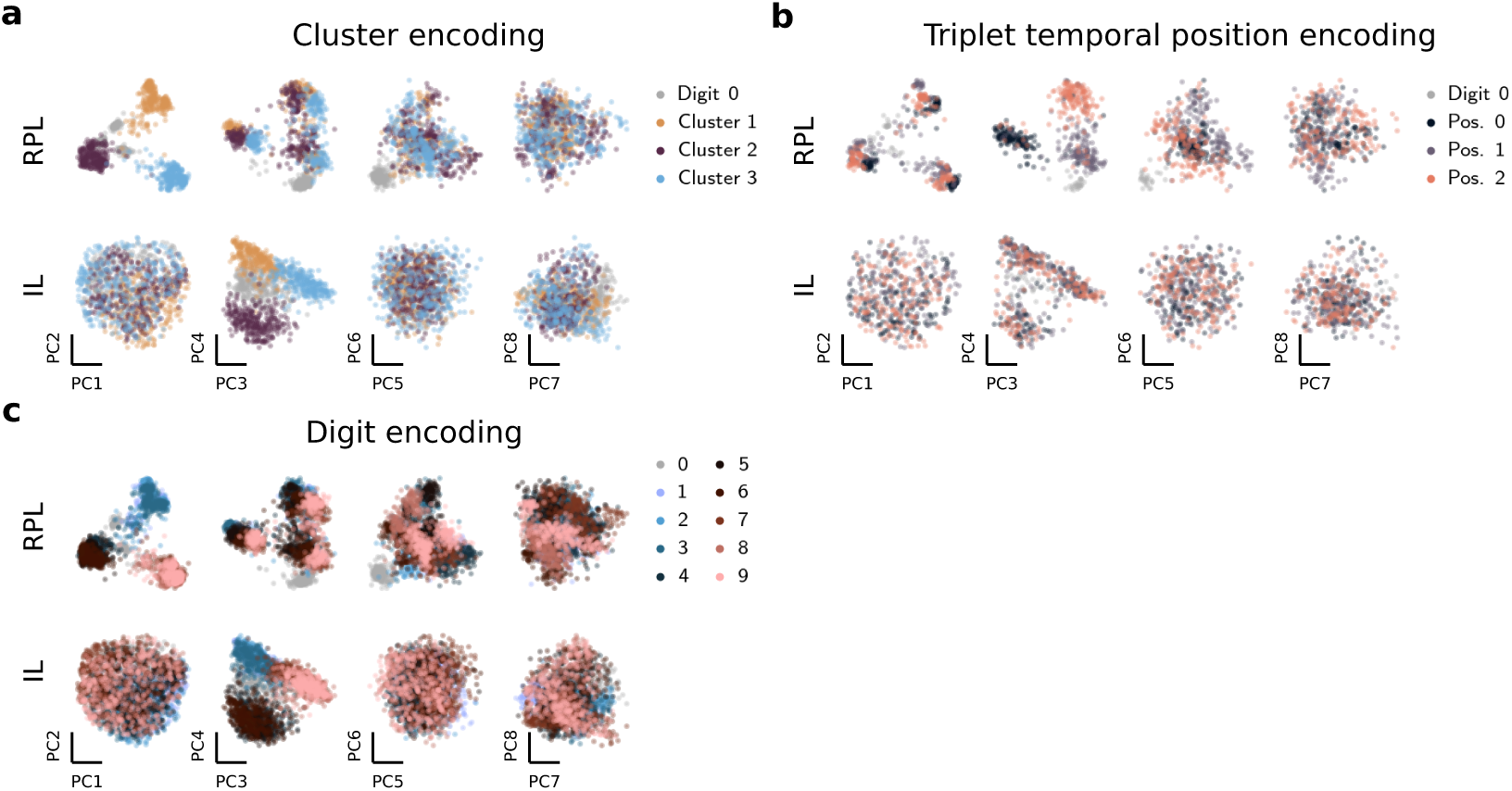
Principal component projections of the learned representations colored by the different latent variables. **(a)** Same as ***Figure 6d*** but for PCs 1–8. **(b)** Same as ***Figure 6e*** but for PCs 1–8. (c) Same as (a) and (b) but colored by digit identity.

**Figure 9—figure supplement 1.**
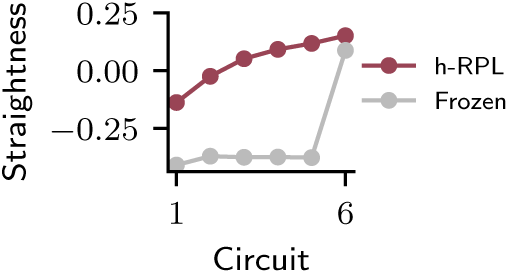
Straightness of learned representations increases over the hierarchy and over learning. The straightness measured as the cosine similarity between consecutive representational difference vectors for h-RPL and the Frozen baseline across the network hierarchy.

**Figure 9—figure supplement 2.**
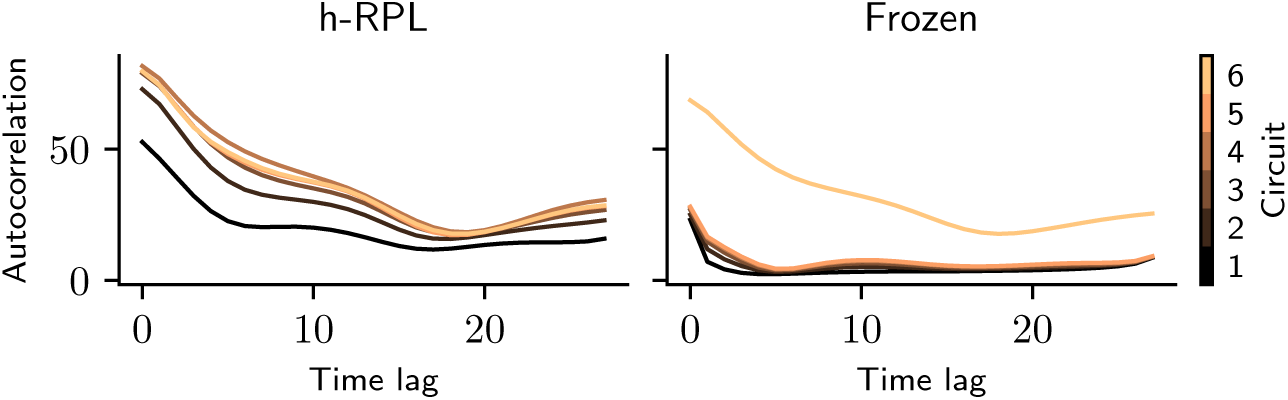
Autocorrelations of learned representations increase over the hierarchy and over learning. The Frobenius norm of the autocorrelation matrix of representations in h-RPL (left) and the frozen control (right) for measuring temporal stability. Autocorrelation increases over the hierarchy for h-RPL and with learning (cf. Frozen vs. h-RPL) across all time lags. The first four frames are excluded from the analysis to reduce the edge effects.

## Footnotes

1 https://github.com/fmi-basel/recurrent-predictive-learning

2 https://www.openslr.org/resources/12/train-clean-100.tar.gz

3 https://drive.google.com/drive/folders/1BhJ2umKH3whguxMwifaKtSra0TgAbtfb

4 https://zenkelab.org/datasets/TLT12

5 https://doi.org/10.5281/zenodo.22141034

6 https://github.com/fzenke/3D_latent_traversal_dataset_generator

7 https://doi.org/10.6084/m9.figshare.19213272.v1

8 https://figshare.com/articles/dataset/Electrophysiological_data_from_the_local-global_experiment/24311767/1

9 https://github.com/mebellet/PFCSequenceModels

