## Supplementary Material for "Understanding neural circuit principles for representation learning through joint-embedding predictive architectures"

#### Supplementary Tables

Supplementary Table S1: Linear decoding performance of learned representations from moving animal videos using different learning objectives (cf. Figure 2–figure supplement 2). "InfoNCE" and "CPC" provide the performance baseline of models trained using a contrastive self-supervised learning (SSL) objective [4] with multi-layer (InfoNCE) and linear predictors (CPC). "Sup." corresponds to a model trained end-to-end using supervised learning. "Rand." corresponds to the performance of a randomly initialized network and "Pixel" corresponds to linear decoding from raw pixel values. All values correspond to mean ± standard deviation ($n = 3$). One model became unstable during training with an $R^2 = -158.77$. We ommitted this value from the average marked with $^*$.

| Variable | $S$ | $\boldsymbol{r}$ | $\boldsymbol{v}$ | $\boldsymbol{\theta}$ | $\omega$ |
|---|---|---|---|---|---|
| Metric | Acc. | $R^2$ | | | |
| RPL | 1.0 | $0.991 \pm 0.001$ | $0.88 \pm 0.01$ | $0.77 \pm 0.03$ | $0.95 \pm 0.04$ |
| CL | $0.99 \pm 0.02$ | $0.997 \pm 0.001$ | $0.908 \pm 0.011$ | $0.4 \pm 0.2$ | $0.012 \pm 0.004$ |
| IL | $0.992 \pm 0.014$ | $0.94 \pm 0.03$ | $0.30 \pm 0.15$ | $0.16 \pm 0.13$ | $0.013 \pm 0.002$ |
| $\text{IL}^c$ | $0.87 \pm 0.01$ | $0.36 \pm 0.16$ | $0.08 \pm 0.01$ | $0.006 \pm 0.001$ | $0.021 \pm 0.005$ |
| $\text{IL}^z$ | $0.862 \pm 0.004$ | $-0.01 \pm 0.02$ | $-0.028 \pm 0.015$ | $-0.01 \pm 0.05$ $^*$ | $0.003 \pm 0.005$ |
| InfoNCE | 1.0 | $0.988 \pm 0.001$ | $0.816 \pm 0.004$ | $0.066 \pm 0.014$ | $0.030 \pm 0.003$ |
| CPC | 1.0 | $0.984 \pm 0.001$ | $0.78 \pm 0.01$ | $0.4 \pm 0.1$ | $0.032 \pm 0.004$ |
| Sup. ($n = 1$) | 1.0 | 0.997 | 0.997 | 0.999 | 1.0 |
| Rand. ($n = 1$) | 0.647 | 0.659 | 0.125 | −0.008 | −0.003 |
| Pixel ($n = 1$) | 0.948 | 0.257 | −0.018 | −0.013 | −3.017 |

Supplementary Table S2: Linear decoding performance of learned representations from Temporal Latent Traversal Twelve (TLT12) dataset using different learning objectives (cf. Figure 4c). "Sup." corresponds to a model trained end-to-end using supervised learning. "Rand." corresponds to the performance of a randomly initialized network. All values correspond to mean ± standard deviation ($n = 5$). We additionally provide the decoding performance on representations learned using the variance-invariance-covariance (VIC) regularization (VICReg) alone for $n = 4$ models and additionally assess the encoder embeddings (VIC (enc)).

| Method | RPL | RPLVIC | IL | $IL^{VIC}$ | VIC | VIC (enc) | Sup. | Rand. |
|---|---|---|---|---|---|---|---|---|
| Category | $0.53 \pm 0.13$ | $0.90 \pm 0.01$ | $0.26 \pm 0.09$ | $0.23 \pm 0.08$ | $0.09 \pm 0.00$ | $0.10 \pm 0.00$ | $1.00 \pm 0.00$ | $0.31 \pm 0.01$ |
| $r_1$ | $0.85 \pm 0.13$ | $0.96 \pm 0.00$ | $0.45 \pm 0.36$ | $0.18 \pm 0.08$ | $0.00 \pm 0.00$ | $0.00 \pm 0.00$ | $0.99 \pm 0.00$ | $0.52 \pm 0.01$ |
| $r_2$ | $0.56 \pm 0.08$ | $0.80 \pm 0.01$ | $0.29 \pm 0.20$ | $0.21 \pm 0.20$ | $0.00 \pm 0.00$ | $0.00 \pm 0.00$ | $0.98 \pm 0.00$ | $0.31 \pm 0.01$ |
| $r_3$ | $0.60 \pm 0.08$ | $0.79 \pm 0.01$ | $0.32 \pm 0.21$ | $0.23 \pm 0.20$ | $0.00 \pm 0.00$ | $0.00 \pm 0.00$ | $0.98 \pm 0.00$ | $0.33 \pm 0.01$ |
| $v_1$ | $0.19 \pm 0.18$ | $0.55 \pm 0.07$ | $0.02 \pm 0.03$ | $0.00 \pm 0.00$ | $0.00 \pm 0.00$ | $0.00 \pm 0.00$ | $0.45 \pm 0.28$ | $0.02 \pm 0.00$ |
| $v_2$ | $0.15 \pm 0.08$ | $0.36 \pm 0.04$ | $0.01 \pm 0.02$ | $0.02 \pm 0.03$ | $0.00 \pm 0.00$ | $0.00 \pm 0.00$ | $0.21 \pm 0.14$ | $0.01 \pm 0.00$ |
| $v_3$ | $0.19 \pm 0.09$ | $0.38 \pm 0.06$ | $0.02 \pm 0.03$ | $0.01 \pm 0.02$ | $0.00 \pm 0.00$ | $0.00 \pm 0.00$ | $0.27 \pm 0.07$ | $0.02 \pm 0.00$ |
| $\theta_1$ | $0.00 \pm 0.00$ | $0.02 \pm 0.00$ | $0.00 \pm 0.00$ | $0.00 \pm 0.00$ | $0.00 \pm 0.00$ | $0.00 \pm 0.00$ | $0.89 \pm 0.00$ | $0.00 \pm 0.00$ |
| $\theta_2$ | $-0.01 \pm 0.00$ | $-0.01 \pm 0.00$ | $0.00 \pm 0.00$ | $0.00 \pm 0.00$ | $0.00 \pm 0.00$ | $0.00 \pm 0.00$ | $0.89 \pm 0.00$ | $-0.01 \pm 0.00$ |
| $\theta_3$ | $0.02 \pm 0.01$ | $0.08 \pm 0.01$ | $0.00 \pm 0.00$ | $0.00 \pm 0.00$ | $-1.67 \pm 0.15$ | $-1.84 \pm 0.10$ | $0.88 \pm 0.00$ | $-0.01 \pm 0.00$ |
| $\omega_1$ | $0.00 \pm 0.00$ | $0.00 \pm 0.00$ | $0.00 \pm 0.00$ | $0.00 \pm 0.00$ | $0.00 \pm 0.00$ | $0.00 \pm 0.00$ | $0.07 \pm 0.01$ | $0.00 \pm 0.00$ |
| $\omega_2$ | $0.00 \pm 0.00$ | $-0.01 \pm 0.00$ | $0.00 \pm 0.00$ | $0.00 \pm 0.00$ | $0.00 \pm 0.00$ | $0.00 \pm 0.00$ | $0.04 \pm 0.00$ | $0.00 \pm 0.00$ |
| $\omega_3$ | $0.00 \pm 0.00$ | $0.01 \pm 0.00$ | $0.00 \pm 0.00$ | $0.00 \pm 0.00$ | $0.00 \pm 0.00$ | $0.00 \pm 0.00$ | $0.31 \pm 0.02$ | $0.00 \pm 0.00$ |
| $\theta^{spot}$ | $0.22 \pm 0.25$ | $0.34 \pm 0.04$ | $0.01 \pm 0.01$ | $0.02 \pm 0.01$ | $0.01 \pm 0.00$ | $0.07 \pm 0.01$ | $0.99 \pm 0.00$ | $0.13 \pm 0.02$ |
| $\phi^{spot}$ | $0.16 \pm 0.16$ | $0.22 \pm 0.04$ | $0.00 \pm 0.01$ | $0.01 \pm 0.00$ | $0.00 \pm 0.00$ | $0.00 \pm 0.00$ | $0.95 \pm 0.00$ | $0.12 \pm 0.01$ |
| Spot hue | $0.54 \pm 0.37$ | $0.77 \pm 0.04$ | $0.20 \pm 0.15$ | $0.27 \pm 0.10$ | $0.62 \pm 0.02$ | $0.72 \pm 0.02$ | $1.00 \pm 0.00$ | $0.56 \pm 0.03$ |
| Floor hue | $0.97 \pm 0.03$ | $0.98 \pm 0.00$ | $0.93 \pm 0.01$ | $0.91 \pm 0.05$ | $0.69 \pm 0.01$ | $0.83 \pm 0.01$ | $1.00 \pm 0.00$ | $0.96 \pm 0.00$ |
| $\Delta\theta^{spot}$ | $0.00 \pm 0.01$ | $0.01 \pm 0.01$ | $0.00 \pm 0.00$ | $0.00 \pm 0.00$ | $0.00 \pm 0.00$ | $0.00 \pm 0.00$ | $0.03 \pm 0.03$ | $0.00 \pm 0.00$ |
| $\Delta\phi^{spot}$ | $0.01 \pm 0.01$ | $0.01 \pm 0.00$ | $0.00 \pm 0.00$ | $0.00 \pm 0.00$ | $0.00 \pm 0.00$ | $0.00 \pm 0.00$ | $0.15 \pm 0.03$ | $0.00 \pm 0.00$ |
| ΔSpot hue | $0.01 \pm 0.01$ | $0.00 \pm 0.00$ | $0.01 \pm 0.00$ | $0.01 \pm 0.00$ | $0.01 \pm 0.00$ | $0.00 \pm 0.00$ | $0.43 \pm 0.01$ | $0.01 \pm 0.00$ |
| ΔFloor hue | $0.36 \pm 0.42$ | $0.14 \pm 0.05$ | $0.01 \pm 0.00$ | $0.04 \pm 0.04$ | $0.00 \pm 0.02$ | $0.00 \pm 0.00$ | $0.46 \pm 0.01$ | $0.07 \pm 0.01$ |

Supplementary Table S3: Linear decoding performance of different models on out-of-distribution test samples from TLT12 dataset (cf. Figure 4–figure supplement 6). "Sup." corresponds to a model trained end-to-end using supervised learning. All values correspond to mean ± standard deviation ($n = 5$).

| Method | RPL | RPLVIC | IL | IL$^{\text{VIC}}$ | Sup. |
|---|---|---|---|---|---|
| Category | $0.41 \pm 0.07$ | $0.54 \pm 0.02$ | $0.21 \pm 0.06$ | $0.19 \pm 0.06$ | $0.84 \pm 0.00$ |
| $r_1$ | $0.87 \pm 0.10$ | $0.97 \pm 0.00$ | $0.47 \pm 0.37$ | $0.19 \pm 0.08$ | $0.99 \pm 0.00$ |
| $r_2$ | $0.57 \pm 0.07$ | $0.78 \pm 0.02$ | $0.30 \pm 0.20$ | $0.22 \pm 0.20$ | $0.97 \pm 0.00$ |
| $r_3$ | $0.63 \pm 0.06$ | $0.77 \pm 0.01$ | $0.33 \pm 0.22$ | $0.24 \pm 0.21$ | $0.95 \pm 0.00$ |
| $v_1$ | $0.21 \pm 0.16$ | $0.53 \pm 0.06$ | $0.02 \pm 0.03$ | $0.00 \pm 0.00$ | $0.43 \pm 0.27$ |
| $v_2$ | $0.16 \pm 0.07$ | $0.35 \pm 0.04$ | $0.01 \pm 0.02$ | $0.02 \pm 0.03$ | $0.20 \pm 0.14$ |
| $v_3$ | $0.20 \pm 0.07$ | $0.37 \pm 0.05$ | $0.02 \pm 0.03$ | $0.01 \pm 0.02$ | $0.24 \pm 0.07$ |
| $\theta_1$ | $-0.01 \pm 0.02$ | $0.00 \pm 0.01$ | $0.00 \pm 0.00$ | $0.00 \pm 0.00$ | $0.14 \pm 0.02$ |
| $\theta_2$ | $-0.01 \pm 0.01$ | $-0.01 \pm 0.00$ | $0.00 \pm 0.00$ | $0.00 \pm 0.00$ | $0.18 \pm 0.04$ |
| $\theta_3$ | $0.01 \pm 0.01$ | $0.04 \pm 0.00$ | $0.00 \pm 0.00$ | $0.00 \pm 0.00$ | $0.27 \pm 0.03$ |
| $\omega_1$ | $0.00 \pm 0.00$ | $0.00 \pm 0.00$ | $0.00 \pm 0.00$ | $0.00 \pm 0.00$ | $-0.02 \pm 0.01$ |
| $\omega_2$ | $-0.01 \pm 0.00$ | $-0.01 \pm 0.00$ | $0.00 \pm 0.00$ | $0.00 \pm 0.00$ | $-0.02 \pm 0.01$ |
| $\omega_3$ | $0.00 \pm 0.00$ | $0.00 \pm 0.00$ | $0.00 \pm 0.00$ | $0.00 \pm 0.00$ | $0.05 \pm 0.01$ |
| $\theta^{\text{spot}}$ | $0.17 \pm 0.23$ | $0.32 \pm 0.05$ | $0.01 \pm 0.01$ | $0.01 \pm 0.01$ | $0.99 \pm 0.00$ |
| $\phi^{\text{spot}}$ | $0.13 \pm 0.15$ | $0.20 \pm 0.04$ | $0.00 \pm 0.01$ | $0.01 \pm 0.01$ | $0.95 \pm 0.00$ |
| Spot hue | $0.44 \pm 0.39$ | $0.74 \pm 0.04$ | $0.19 \pm 0.15$ | $0.27 \pm 0.10$ | $1.00 \pm 0.00$ |
| Floor hue | $0.96 \pm 0.04$ | $0.98 \pm 0.00$ | $0.93 \pm 0.01$ | $0.92 \pm 0.05$ | $1.00 \pm 0.00$ |
| $\Delta\theta^{\text{spot}}$ | $0.00 \pm 0.01$ | $0.01 \pm 0.01$ | $0.00 \pm 0.00$ | $0.00 \pm 0.00$ | $0.03 \pm 0.03$ |
| $\Delta\phi^{\text{spot}}$ | $0.00 \pm 0.01$ | $0.01 \pm 0.00$ | $0.00 \pm 0.00$ | $0.00 \pm 0.00$ | $0.15 \pm 0.03$ |
| ΔSpot hue | $0.01 \pm 0.01$ | $0.00 \pm 0.00$ | $0.01 \pm 0.00$ | $0.01 \pm 0.00$ | $0.42 \pm 0.02$ |
| ΔFloor hue | $0.29 \pm 0.40$ | $0.14 \pm 0.05$ | $0.01 \pm 0.00$ | $0.04 \pm 0.04$ | $0.45 \pm 0.01$ |

Supplementary Table S4: Linear regression $R^2$ (mean ± std.) of representations learned from mouse in open arena videos with different learning objectives (cf. Figure 5b). "Sup." represent the performance of a model trained with ground-truth labels. "Rand." provides the baseline performance of a randomly initialized network (see Methods).

| Task | $\boldsymbol{r}_N$ | $\boldsymbol{v}_T$ | $\boldsymbol{\theta}$ |
|---|---|---|---|
| RPL | $0.67 \pm 0.02$ | $0.08 \pm 0.02$ | $0.11 \pm 0.01$ |
| IL | $0.21 \pm 0.11$ | $0.04 \pm 0.01$ | $0.03 \pm 0.02$ |
| Sup. ($n = 1$) | 0.65 | 0.2 | 0.27 |
| Rand. ($n = 1$) | 0.54 | 0.0 | −0.01 |

Supplementary Table S5: Linear classification accuracy (mean ± std.) of representations learned from natural speech recordings with different learning objectives (cf. Figure 5d). "Sup." represent the performance of models trained with ground-truth labels optimized for each task. "Rand." provides the baseline performance of a randomly initialized network (see Methods).

| Task | Speaker | Phoneme |
|---|---|---|
| RPL | $0.982 \pm 0.002$ | $0.600 \pm 0.001$ |
| IL | $0.96 \pm 0.01$ | $0.26 \pm 0.01$ |
| Sup. ($n = 1$) | 0.99 | 0.65 |
| Rand. ($n = 1$) | 0.54 | 0.31 |

Supplementary Table S6: Linear classification accuracy (mean ± std.) of representations learned from MNIST digit sequences with different learning objectives (cf. Figure 6d). "Sup." represent the performance of a model trained with ground-truth labels. "Rand." provides the baseline performance of a randomly initialized network (see Methods).

| Task | Cluster | Digit | Triplet |
|---|---|---|---|
| RPL | $0.994 \pm 0.001$ | $0.978 \pm 0.003$ | $0.965 \pm 0.006$ |
| IL | $0.979 \pm 0.006$ | $0.69 \pm 0.11$ | $0.44 \pm 0.15$ |
| Sup. ($n = 1$) | 0.994 | 0.983 | 0.983 |
| Rand. ($n = 1$) | 0.778 | 0.663 | 0.608 |

### Supplementary Notes

#### S1 The role of predictor capacity in learned representations in recurrent predictive learning (RPL)

Prior machine learning studies have shown that joint-embedding predictive architectures (JEPAs) with higher predictor capacity lead to better downstream performance on the learned representations from visual stimuli [2, 1, 3]. We wondered if similar effects can be observed in RPL. For this purpose, we trained RPL models with different nonlinear predictor architectures on the moving animals videos, where we used Multi-Layer Perceptrons (MLPs) with different widths (i.e., one hidden layer with varying number of hidden units) and depths (i.e., varying number of hidden layers, each with 512 units). To evaluate downstream performance as a function of predictor's width and depth, we followed the same evaluation as before (Methods) and then compute the average $R^2$ score over all dynamic variables. While the lower predictor width resulted in quantitatively worse representations, the predictor's depth had no effect on downstream performance (Supplementary Fig. S1).

It is worth to mention that the integrator in RPL directly influences the prediction computation; hence its capacity will influence the learned representations from a JEPA. However, this influence in the downstream performance is convoluted with the effect of representation dimensionality and does not lead to concrete understanding of computational capacity of the prediction.

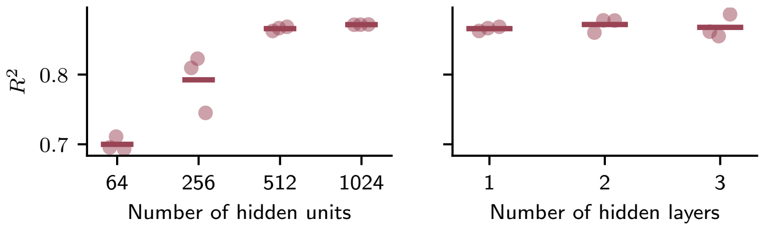


Supplementary Figure S1: **Downstream performance as a function of predictor's capacity in RPL models.** Average $R^2$ score over all regression tasks in the moving animals dataset as a function of number of hidden units (left) and number of hidden layers (right) in the nonlinear predictor in RPL models.

#### S2 Encoder representations on the MNIST sequence task

To check whether RPL also results in higher decoding accuracy than invariance learning (IL) when evaluated on the embeddings $\boldsymbol{z}$, we trained three classifiers on the learned embeddings $\boldsymbol{z}$ from the MNIST digit triplet sequences for cluster identity, digit identity, and triplet identity tasks (cf. Figure 6a-c). In contrast to representations $\boldsymbol{c}$, the embeddings resulted in high and low accuracy for cluster and triplet tasks, respectively, for both RPL and IL (Supplementary Fig. S2a,b). This was expected because the encoder does not have access to the temporal context of the sequence which is essential for the triplet classification. Additionally, we observed that IL obtained lower accuracy than RPL, but still higher than a randomly initialized network, for digit classification task. This finding seems to be at odds with the accuracy levels obtained by decoding from the representations $\boldsymbol{c}$ for IL (Supplementary Fig. S2a and cf. Figure 6c). We hypothesized that the underlying reason for this discrepancy is that the recurrent representations $\boldsymbol{c}$ are more directly optimized for the invariance objective and therefore collapse the digits from the same cluster together; while the feedforward encoder, could still retain some information about the individual MNIST digits. We validated this hypothesis by training a similar network, but without an integrator component, and found that the task accuracies from the embeddings $\boldsymbol{z}$ in this modified network now matched the performance obtained from representations $\boldsymbol{c}$ in full RPL networks (Supplementary Fig. S2a,c).

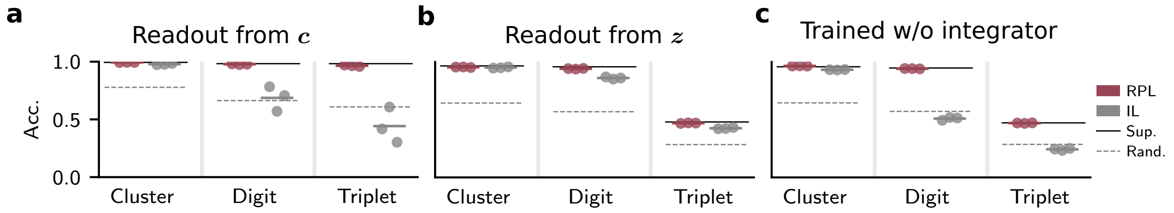


Supplementary Figure S2: **Readout accuracy from integrator and encoder on the abstract MNIST sequence task.** **(a)** Same as Figure 6c. **(b)** Same as (a) but for the encoder. **(c)** Same as (b) but for the network trained without an integrator.

#### S3 Phase-corrected rotation probe as a bilinear optimization problem

To probe whether the learned representation $\mathbf{c}$ encodes object orientation $\theta_3$, we fit a linear decoder from $\mathbf{c}$ to the ground-truth rotation angle $\theta = \theta_3 \in [-\pi, \pi]$ about the $z$-axis. Because different object categories have unknown canonical orientations, a single global decoder trained on the targets $[\sin\theta, \cos\theta]^\top$ would conflate this per-category offset with the rotation signal and systematically underestimate decodability. We therefore adopt a probe model with a *shared* linear decoder $\mathbf{W} \in \mathbb{R}^{2\times d_c}$ and *per-category* phase offsets $\{\phi_k\}_{k=1}^{K}$:

$$\mathbf{W}\mathbf{c}_i^{(k)} \approx \begin{bmatrix} \sin\big(\theta_i^{(k)} + \phi_k\big) \\ \cos\big(\theta_i^{(k)} + \phi_k\big) \end{bmatrix}. \tag{1}$$

The shared $\mathbf{W}$ enforces the hypothesis that all categories rely on the same two-dimensional linear subspace of $\mathbf{c}$ to represent rotation, while $\phi_k$ absorbs the unknown canonical orientation of each category.

**Objective.** Let $\mathbf{C}_k \in \mathbb{R}^{n_k\times d_c}$ be the matrix of training representations for category $k$, and let $\mathbf{T}_k \in \mathbb{R}^{n_k\times 2}$ contain the unphased targets $[\sin\theta_i^{(k)}, \cos\theta_i^{(k)}]$ as rows. Using the angle-addition identity

$$\begin{bmatrix} \sin(\theta+\phi) \\ \cos(\theta+\phi) \end{bmatrix} = \mathbf{R}(\phi) \begin{bmatrix} \sin\theta \\ \cos\theta \end{bmatrix}, \qquad \mathbf{R}(\phi) := \begin{bmatrix} \cos\phi & -\sin\phi \\ \sin\phi & \cos\phi \end{bmatrix}, \tag{2}$$

the phased target matrix for category $k$ is $\mathbf{T}_k(\phi_k) = \mathbf{T}_k\,\mathbf{R}(\phi_k)$. We minimize the $\ell_2$-regularized objective

$$\mathcal{L}(\mathbf{W}, \{\phi_k\}) = \sum_{k=1}^{K} \big\|\mathbf{C}_k\mathbf{W}^\top - \mathbf{T}_k(\phi_k)\big\|_F^2 + \alpha\,\|\mathbf{W}\|_F^2, \tag{3}$$

where $\alpha > 0$ is the ridge regularization strength. The problem is bilinear—and therefore non-convex—in $(\mathbf{W}, \{\phi_k\})$ jointly, but becomes convex in each block when the other is held fixed.

**Alternating minimization.** We exploit this block-convex structure and alternate two closed-form updates.

*Step A — phase update.* Given $\mathbf{W}$, the cost for category $k$ depends on $\phi_k$ only through $\mathrm{tr}(\mathbf{A}_k\,\mathbf{R}(\phi_k))$, where $\mathbf{A}_k = \hat{\mathbf{Y}}_k^\top \mathbf{T}_k \in \mathbb{R}^{2\times 2}$ and $\hat{\mathbf{Y}}_k = \mathbf{C}_k\mathbf{W}^\top$ are the current predictions. Expanding the trace gives

$$\mathrm{tr}(\mathbf{A}_k\,\mathbf{R}(\phi_k)) = (A_{00} + A_{11})\cos\phi_k + (A_{01} - A_{10})\sin\phi_k, \tag{4}$$

which is maximized in closed form by

$$\phi_k^* = \mathrm{atan2}(A_{01} - A_{10},\ A_{00} + A_{11})\,. \tag{5}$$

This step scales as $\mathcal{O}(n_k d_c)$ per category and introduces no additional hyper-parameters.

*Step B — decoder update.* Given $\{\phi_k\}$, define the rotated targets $\tilde{\mathbf{T}}_k = \mathbf{T}_k(\phi_k)$ and concatenate across classes: $\mathbf{C} = [\mathbf{C}_1; \ldots; \mathbf{C}_K]$, $\tilde{\mathbf{T}} = [\tilde{\mathbf{T}}_1; \ldots; \tilde{\mathbf{T}}_K]$. The update reduces to standard ridge regression,

$$\mathbf{W}^* = \tilde{\mathbf{T}}^\top \mathbf{C}\left(\mathbf{C}^\top\mathbf{C} + \alpha\mathbf{I}\right)^{-1}, \tag{6}$$

which is solved efficiently via the normal equations.

**Initialization and convergence.** We initialize $\mathbf{W}$ by running Step B with all $\phi_k = 0$ (i.e. plain global ridge regression on the unphased targets), then alternate Steps A and B for 20 iterations. The objective in Eq. (3) decreases monotonically because each step minimizes over its own block. In practice convergence is reached in fewer than ten iterations.

**Evaluation.** After fitting, we freeze $\mathbf{W}$ and $\{\phi_k\}$ and compute $R^2$ between the predicted and ground-truth targets on the held-out test set. Random-initialization baselines use representations from an untrained network evaluated with the same procedure.

[4] A. v. d. Oord, Y. Li, and O. Vinyals. Representation Learning with Contrastive Predictive Coding. *arXiv:1807.03748 [cs, stat]*, 2018. arXiv: 1807.03748.
